# Haplotype-resolved genome of autotetraploid alfalfa (*Medicago sativa*) Regen-SY27x uncovers large scale structural variation and resistance gene dynamics

**DOI:** 10.64898/2026.05.01.722254

**Authors:** Harpreet Kaur, Cameron Connor, Adam Gomez, Joann Mudge, Andrew Farmer, Laura M. Shannon, Deborah A. Samac

## Abstract

Polyploid genome assembly presents unique challenges due to extensive heterozygosity and complex haplotype structure. We report a haplotype-resolved, chromosome-scale assembly of Regen-SY27x, a genotype of autotetraploid alfalfa (*Medicago sativa*), which is widely used for genetic modification because of its excellent regenerative capacity in tissue culture. Using PacBio HiFi long reads, Omni-C scaffolding, and linkage map guided phasing, we generated a 3.2 GB assembly comprising four haplotypes with high contiguity and completeness. Kmer-based validation confirmed accurate haplotype separation, while linkage map integration and dotplot analysis identified and corrected chimeric scaffolds. Gene annotation yielded 221,688 protein-coding genes, with more than 99% assigned to pseudochromosomes. Repetitive elements accounted for 62.7% of the genome, dominated by long terminal repeat retrotransposons and a high fraction of Helitrons. The spatial enrichment of Helitrons within gene-dense distal chromosome arms underscores their pivotal role as key drivers of genomic innovation and gene family expansion. We identified 3,696 nucleotide-binding leucine-rich repeat R genes, with Toll/interleukin-1 receptor-like and Rx-type subclasses forming large tandem clusters across haplotypes. Comparative analyses revealed strong macrosyntenic conservation among Regen-SY27x and the publicly available Chinese alfalfa genomes but extensive structural variation both within Regen-SY27x haplotypes and between Regen-SY27x and the Chinese genotypes with tens of thousands of duplications, inversions, and translocations detected. These results demonstrate that a single autotetraploid individual captures extensive structural diversity, but individuals from different populations vary greatly. The Regen-SY27x assembly provides a foundational genomic resource for investigating polyploid genome evolution and identifying genetic variation relevant to biological and agronomic improvement in alfalfa.

**Article Summary:** This study presents the first chromosome-scale, haplotype-resolved genome assembly of the US alfalfa germplasm, Regen-SY27x, a key alfalfa genotype used widely for genetic engineering. We integrated HiFi long reads, Omni-C^TM^ scaffolding, and linkage map-guided phasing to reconstruct all four haplotypes of this complex autotetraploid. Our results identified 221,688 protein-coding genes and reveal immense intra-individual structural variations dominated by small duplications. This high-quality reference serves as a foundational tool for the alfalfa community, enabling researchers to link complex structural diversity with agronomic traits and further enhance the biotechnological potential of this essential forage crop.

## Introduction

High-quality crop plant genome assemblies are crucial for both facilitating crop improvement and understanding traits, genome structure, and genetic variation. Sequencing technologies have evolved significantly over the last few decades, from Sanger sequencing in the late 1970s to next-generation sequencing (NGS) platforms like Illumina HiSeq and MiSeq, and more recently, third-generation sequencing technologies such as Pacific Biosciences High-Fidelity (PacBio HiFi) and Oxford Nanopore Technology (ONT). These advancements, coupled with progress in bioinformatics, have enhanced sequencing speed, accuracy, and cost-effectiveness (Jiao and Schneeberger 2017; Marks et al. 2021). Polyploid species, especially autopolyploids which do not exhibit preferential pairing at meiosis, present challenges in genome assembly compared to diploids due to the presence of multiple copies of each chromosome, making it more difficult to accurately resolve and assemble homologous sequences (Wang et al. 2023). The complexity escalates with greater heterozygosity, increasing challenges in distinguishing allelic variation from paralogous sequences, which may result in fragmented assemblies. Additionally, the larger genome size and higher repetitive element content commonly found in polyploids makes assembling a genome even more challenging (Kyriakidou et al. 2018; Kyriakidou et al. 2020). Efforts have been undertaken to address these hurdles using long-read sequencing along with long-range scaffolding techniques such as optical mapping (Belser et al. 2018; Pan and Lonardi 2019) and high throughput chromosome conformation capture (Burton et al. 2013; Zhang et al. 2019).

Alfalfa (*Medicago sativa* L.), hailed as the “Queen of Forages,” is a valuable perennial forage legume in the United States, accounting for a production value of $9.52 billion in 2024, the fourth highest following corn, wheat, and soybean (United States Department of Agriculture 2025). It is a highly heterozygous, cross pollinated, autotetraploid species. Alfalfa plays important functions within sustainable cropping systems (Sheaffer and Seguin 2003). The primary use of alfalfa in the United States is as a forage crop for livestock feed, particularly for dairy cows, beef cattle, horses, and sheep (Conrad and Klopfenstein 1988). The high protein content, digestibility, and nutrient profile of alfalfa make it an essential component of many livestock diets, contributing to the health, growth, and productivity of animals (Coburn, Wells, Sheaffer, et al. 2021). Alfalfa can be utilized in various other ways, such as increasing soil fertility through symbiotic nitrogen fixation (Sheaffer and Seguin 2003), contributing to soil carbon and nitrogen retention (King and Hofmockel 2017), aiding in environmental cleanup through phytoremediation (Chekol and Vough 2001; Miller et al. 2022; Tussipkan and Manabayeva 2022), feeding fish (Coburn, Wells, Phelps, et al. 2021), serving as nutritious sprouts for human consumption, and medicinal applications (Raeeszadeh et al. 2022). Alfalfa additionally offers a habitat for wildlife and acts as a nectar supply for pollinating insects (Fernandez et al. 2019). It can also be used as biofuel feedstock (Sheaffer et al. 2000).

Assembly of genome sequences of tetraploid alfalfa is complicated by strong inbreeding depression preventing development of inbred lines. Alfalfa cultivars are bred as synthetic populations of heterogenous outcrossing individuals. Thus, published genome sequences of alfalfa have been assembled from DNA of a single genotype selected from a named cultivar. Initially, the alfalfa community depended on the diploid *M. truncatula* (Tang et al. 2014; Pecrix et al. 2018) reference genome for genomic analyses due to its high synteny with *M. sativa* (Julier et al. 2003; Choi et al. 2004; Li, Wei, et al. 2014; Ray et al. 2015). Early attempts in 2013-2015 were made to sequence a tetraploid alfalfa genotype, NECS-141, selected from an experimental breeding population adapted to the midwestern United States, but the data remained unpublished (Khu et al. 2013; Mudge and Farmer 2021). Two diploid accessions of *M. sativa* from the U.S. germplasm have been sequenced and assembled: cultivated alfalfa at the diploid level (CADL) with 5,753 contigs (https://data.legumeinfo.org/Medicago/sativa/genomes/), and *Medicago sativa* spp. *caerulea* PI 464715at the chromosome scale (Li et al. 2020). Recently, three alfalfa genome assemblies from Chinese cultivars have been published: the consensus chromosome-scale Zhongmu No. 1 (Shen et al. 2020), and the haplotype-aware chromosome-scale XinJiangDaYe (Chen et al. 2020) and Zhongmu No. 4 (Long et al. 2022) hereafter abbreviated Zm1, XJDY, and Zm4, respectively. For thousands of years, China has grown alfalfa with minimal external influence (Russelle 2001). This has resulted in distinct genetic traits unique to Chinese alfalfa (Wang and Şakiroğlu 2021). Therefore, it is beneficial to sequence an alfalfa genome using germplasm from outside China, as alfalfa germplasm from other regions possesses unique traits.

In this study, we generated a haplotype-resolved, chromosome-scale assembly of the autotetraploid alfalfa genotype Regen-SY27x, representing North American germplasm by integrating PacBio HiFi sequencing, Omni-C scaffolding, and linkage map–guided phasing. We then assessed assembly quality, annotated genes, repeats, and resistance gene families, and examined the contribution of transposable elements to genome organization. Finally, comparative analyses within Regen-SY27x and across the sequenced Chinese genotypes revealed patterns of structural variation and gene family dynamics, providing new insights into polyploid genome evolution and resources for alfalfa improvement.

## Materials and methods

### Plant material

A single genotype, Regen-SY27x, was selected for genome sequencing and assembling. This plant possesses high regenerative ability under tissue culture conditions and has been widely used in plant transformation experiments (Samac and Austin-Phillips 2006; Samac and Temple 2021). Regen-SY27x was selected from the cultivar Regen-SY (Bingham 1991). Regen-SY is an F_1_ hybrid between first generation selfed (S_1_) progenies of Regen-S (*M. sativa*) (Bingham 1989) and Regen-Y (*M. falcata*) (Bingham et al. 1975) which were developed using recurrent selection for high regenerative ability under *in vitro* conditions.

The Regen-SY27x genotype was used as one of the parents to develop a biparental F_1_ mapping population to construct genetic maps. Single plants for each of the two parents, Regen-SY27x and an individual plant from MN Bio I C3 created as a bioenergy alfalfa for non-lodging stems (Lamb et al. 2014), were intermated by hand pollination without emasculation to generate 165 F_1_ progenies. The plants were grown and maintained in a greenhouse using a potting mix (Metro-Mix, Sun Gro Horticulture) at 24°C day/night temperatures with a photoperiod of 16 h.

### DNA and RNA isolation

#### Genomic DNA for PacBio sequencing

Immature leaves that had been held in darkness for 48 h were removed from the plant, immediately frozen in liquid nitrogen and stored at −80°C. The high molecular weight (HMW) genomic DNA extraction and purification from samples was performed using MagAttract HMW DNA Kit (Qiagen Inc., Venlo, Netherlands) as per the manufacturer’s instructions and confirmed using a FemtoPulse (Advanced Analytical Technologies Inc., Ankeny, IA).

#### Genomic DNA for genotyping-by-sequencing

For linkage mapping, the genomic DNA was extracted from leaves of the two parents and 165 F_1_ progenies from 20-week-old plants using Qiagen DNeasy plant kit based on the manufacturer’s protocol. Genotyping-by-sequencing (GBS) for the population on the Illumina NovaSeq^TM^ 6000 (Illumina, CA, USA) using single end 1×100 reads was conducted through the University of Minnesota Genomics Center (https://genomics.umn.edu/services/gbs) using the *Ape*KI single enzyme digestion protocol (See Supplementary Methods for more details).

#### RNA extraction, Illumina RNA-seq and PacBio IsoSeq

The RNA was extracted from 20-week-old leaf, stem, flower, seedpod, root, and nodule tissues. RNA-seq and PacBio Iso-seq data were obtained from these tissues following the protocol outlined in Miller et al. (2022). RNA-seq was performed by GeneWiz (Azenta Life Sciences) and IsoSeq was performed by Seqmantic (Fremont, CA, USA).

### Genome sequencing

#### PacBio sequencing

Genome sequencing was performed by the University of Delaware, Delaware Biotechnology Institute (Newark, DE USA). The 10 µg aliquots of HMW DNA were converted to SMRTbell templates using the HiFi Express Template Prep Kit 2.0 (Pacific Biosciences, Menlo Park, CA) as per the manufacturer’s instructions. In summary, samples were end-repaired and ligated to blunt adaptors. Exonuclease treatment was performed to remove unligated adapters and damaged DNA fragments. Samples were purified using 0.6x AMPureXP beads (Beckman Coulter Inc., Brea, CA). The purified SMRTbell libraries were eluted in 10 μl elution buffer. Eluted SMRTbell libraries were selected based on size using the BluePippin (Sage Science Inc., Beverly, MA) to eliminate library fragments below approximately 10 Kbp. Final library quantification and sizing was carried out on a FemtoPulse (Advanced Analytical Technologies Inc., Ankeny, IA) using 1 μl of library. The amount of primer and polymerase required for the binding reaction was determined using the SMRTbell concentration and library insert size. Primers were annealed and polymerase was bound to SMRTbell templates using the Sequel II binding kit 2.0 (Pacific Biosciences, Menlo Park, CA). Sequencing was performed using the Sequel IIe (Pacific Biosciences, Menlo Park, CA). Sequel IIe loading efficiency was optimized for each individual library utilizing a standardized titration protocol. The HiFi libraries were run on Sequel IIe system 8M SMRT cells using sequencing chemistry 3.0 with 4 -hour pre-extension and 30-hour movie time.

#### Dovetail Omni-C^TM^ mapping

Immature leaves that had been dark treated for 48 h and flash frozen in liquid nitrogen were used for DNA extraction. Omni-C library preparation, quality control, and sequencing was performed by Dovetail Genomics, Scotts Valley, CA, USA.

### Development of SNP markers and linkage maps

SNP markers and linkage maps were generated following the method described by (Kaur et al. 2026). Briefly, SNP markers were developed from GBS raw reads processed with standard quality control using Fastqc v. 0.11.9 and trimming with Trimmomatic software v 0.39 (Bolger et al. 2014). These reads were aligned to the ZhongmuNo.1 (Shen et al. 2020) reference genome and SNPs were called using Next Generation Sequencing Experience Platform (NGSEP) MultisampleVariantCaller v. 4.2.1 (Perea et al. 2016) followed by stringent filtering resulting in SNP markers with an average mean read depth of 78.2 per F_1_ individual (See Supplementary Methods). For linkage map construction, genotypes were called with updog (Gerard et al. 2018), along with additional SNP filtering to keep SNPs with missing proportion < 0.05 and estimated allele bias between 0.2 and 4.5, followed by map construction with Mappoly (Mollinari and Garcia 2019) to account for the autotetraploid nature of alfalfa. The markers were then grouped and ordered using reference genome sequence information and phased to map 2,482 SNPs on 8 linkage groups (Supplementary Table 1). These phased linkage maps were subsequently used for phasing the Regen-SY27x genome assembly. See Supplemental Methods for full details.

### K-mer frequency distribution

Pacbio HiFi data was k-merized using jellyfish 2 (Marçais and Kingsford 2011) at k=27 and plotted with R (4.4) to visualize the k-mer frequency distribution. Peaks were identified to assess the likelihood of assembling all four haplotypes and ensure that the error fraction and the peaks were not overlapping significantly. Read coverage was determined from the leftmost single-copy peak.

### Phased tetraploid genome assembly

PacBio HiFi data was assembled using Hifiasm (0.19.8-r603). Hifiasm was parameterized to create four haploid assemblies of approximately 800 Mb, using Omni-C^TM^ sequence data (--h1 read1 and --h2 read2) to assist in solving the graph. The assembly maximized kmer length (-k) and minimizer window size (-w) at 63 to get more discriminate seeding and increased k-mer occurrence (-D) to 8. In addition, the estimated haploid genome size (--hg-size) was set to 800m, the number of haplotypes (--n-hap) was set to 4, and the --primary flag was used. Deduplication was iteratively performed to determine the best result. As a final step, additional scaffolding was done using SALSA (Ghurye et al. 2017) after following the Omni-C^TM^ best practices for data preparation.

### Haplotype phased pseudomolecule construction using the genetic map

Additional haplotype partitioning was done to improve scaffold placement within phased haplotypes (a set of chromosomes representing a monoploid complement; hereafter referred to as “haplotype”). Whole genome alignments of the scaffolds to ZhongmuNo.1 as the reference were made using minimap2 (Li 2018), and these alignments were used with samtools mpileup (Li et al. 2009) to cross-reference alleles phased in the linkage groups with the occurrence of those same alleles in the genome assembly. This enabled comparison of dosages of markers derived from the GBS data with the assembly and association of scaffolds containing informative single-dosage markers (either non-reference allele for AAAB/1/simplex, or reference allele for ABBB/3/triplex) with the corresponding linkage groups. Assembly haplotypes were broken into chromosomes and associated with one of the 32 linkage groups, maximizing markers shared with the genetic map. As a first pass, scaffolds with at least three markers pointing to the same linkage group were considered anchored to that linkage group and, if necessary, were moved to the matching haplotype. Chimeric scaffolds that had markers in different regions pointing to different linkage groups were broken and moved to the appropriate haplotypes. Visual confirmation using dotplots aligned to PI464715 (Li et al. 2020) using dotplotly (https://github.com/tpoorten/dotPlotly) allowed us to better pinpoint the breakpoints, minimizing overlap with adjacent scaffolds in the same haplotype. At this point, scaffolds with less than three markers anchoring them to the genetic map were moved after checking marker alignments. Because just over half of the nucleotides in the assembly were in scaffolds not anchored to the genetic map, we took additional steps to identify and move scaffolds that appeared to be in the wrong haplotype. We used minimum tiling paths visualized on dotplots and runs of BUSCO genes (see below) that were duplicated within a haplotype. These duplicate scaffolds were moved to other haplotypes if they filled in a gap in the minimum tiling path. The final set of scaffolds, partitioned into 32 chromosome haplotypes were stitched into pseudomolecules with 100Ns between them, ordering and orienting scaffolds based on their position in the genome assembly of *M. sativa* spp. *caerulea* PI464715 (Li et al. 2020). Scaffolds that were not placed into a pseudomolecule were left as individual sequences.

### Assembly quality

The quality of the assembly was assessed using several methods. We calculated basic assembly statistics and ran Benchmarking Universal Single-Copy Orthologs (BUSCO 5.6, Fabales odb10; (Seppey et al. 2019)) on the assembly. The BUSCO completeness and copy number were considered to ensure that the BUSCOs for Fabales were well covered and that most of the BUSCOs were found at the expectation of four copies. In addition, the k-mer analysis toolkit (KAT; (Mapleson et al. 2017)) was used to ensure that the assemblies covered the HiFi read k-mer frequency distribution and further, that the copy numbers seen under the peaks for the assembly matched their expected copy number. Dotplots against *M. sativa* spp. *caerulea* PI464715 were also used to visually inspect the scaffolding order and orientation of the phased assembly before building pseudochromosomes.

### RNA-seq and PacBio Iso-seq analyses

To incorporate gene expression data, a reference-based assembly was generated from the Illumina RNA-seq and PacBio IsoSeq reads as follows. Illumina reads were aligned to each haplotype from the genome assembly using hisat2 v2.2.1 (Kim et al. 2019) using default parameters except the maximum intron length (--max-intronlen) was set to 25000 and --dta, which outputs alignments for downstream transcription assembly, allowing up to five distinct, primary alignments. PacBio IsoSeq reads were aligned to each haplotype of the genome assembly with minimap2 using the splice preset (-x splice) and reducing the max intron length (-G) to 25,000 nts. StringTie (Pertea et al. 2015) was then used to do a reference-based assembly of the transcriptome reads and gffread (Pertea and Pertea 2020) was used to convert the resulting GTF to GFF3 format.

### Gene identification and annotation

Structural annotations were predicted using MAKER (v 3.01.03) (Campbell et al. 2014) following a similar strategy to that outlined in Card et al. (Card et al. 2019). TEtools (https://github.com/Dfam-consortium/TETools) was used to run RepeatModeler2 (Flynn et al. 2020) in order to create a database of repeats from the genome assembly. Repeatmasker (version 4.1.5) was run with the ncbi engine (-e). The complex repeats were separated out for use in MAKER and converted into GFF3 format. The first MAKER round was run passing in the StringTie GFF3 file and the Repeatmasker file GFF3 file. In addition, existing proteome models from *Glycine max* Williams 82 (version 4; (Valliyodan et al. 2019)), *M. truncatula* Jemalong A17 (version 5; (Pecrix et al. 2018)), *M. truncatula* A17_HM341 (version 4, (Tang et al. 2014)), and uniprot Fabaceae proteins (UniProt Consortium 2023) were used. A second MAKER round was run to improve the gene calls. This round used ESTs, proteins, and repeats created in the first MAKER run. In addition, a SNAP (Korf 2004) model, previously created in *Cicer arietinum*, was also used and an Augustus model trained on the assembly using BUSCO’s built-in training procedure (parameters --long --augustus).

### Adjudication of structural annotations

To solve the issue of overlapping gene models from multiple evidence sets, a custom tool called the Adjudicator was developed by NCGR (https://github.com/ncgr/Adjudicator). The script takes sets of structural predictions in GFF3 whose derived proteins have been scored against gene family Hidden Markov Models from the Legume Information System at https://data.legumeinfo.org/LEGUMES/Fabaceae/genefamilies/legume.fam1.M65K (Stai et al. 2019) and creates an interval tree to find and compare overlapping gene predictions. The overlapping predictions are compared, and if they have the same gene family, the model with the better score (within 5% relative by default) is selected. If they do not have the same family, they are compared to other overlapping models. After all of the evidence sets have been adjudicated, the final models and the repeat models are overlapped with BedTools intersect (-wao). The resulting overlaps were filtered to remove predictions with greater than 40% repeat coverage over all exons. As a final step, an analysis was conducted to examine overlapping predictions that could not be solved using the tool above at the CDS level. BUSCO was used to ensure that the resultant collapse did not markedly diminish assembly completeness or eliminate gene models originally detected at a copy number of four.

### Identification and annotation of repeated elements

Repetitive elements in the Regen-SY27x genome were identified and annotated using the Extensive de novo TE Annotator (EDTA v2.2.2) pipeline (Ou et al. 2019). The EDTA workflow integrates multiple transposable element (TE) detection tools and reference libraries to provide a comprehensive annotation of both class I (retrotransposons) and class II (DNA transposons) using various tools. Briefly, the assembled genome was used as input for EDTA v2.2.2. The pipeline first performed *de novo* identification of long terminal repeat (LTR) retrotransposons using LTRharvest (Ellinghaus et al. 2008) and LTR_FINDER (Xu and Wang 2007), followed by structural refinement with LTR_retriever (Ou and Jiang 2018). Non-LTR retrotransposons and DNA transposons were detected with RepeatModeler v2.0.6 (Flynn et al. 2020), while miniature inverted-repeat transposable elements (MITEs) and Helitrons were predicted using MITE-Hunter (Han and Wessler 2010) and HelitronScanner (Xiong et al. 2014), respectively. The outputs were filtered to reduce redundancy and false positives, and the curated library was then used for genome-wide annotation with RepeatMasker (Tempel 2012), producing a final, non-redundant catalog of repeats. The curated TE library was generated using annotated repeats of nine different legume species including *Arachis hypogeae, Vicia faba, M. truncatula, Lotus japonicus, Cajanus cajan, Glycine max, Phaseolus vulgaris, Vigna radiata,* and *Cicer arietinum* obtained from Plantrep database (Luo et al. 2022). The proportions of each repeat type, total repeat content, and distribution across the genome were summarized from the RepeatMasker output.

### Identification of disease resistance (R) genes

Annotation of R genes is challenging due to their frequent overlap with repetitive sequences. Therefore, *de novo* annotation of R genes was conducted for the unmasked Regen-SY27x genome using the FindPlantNLRs pipeline (Chen et al. 2022) using default settings and visualized with RIdeogram (Hao et al. 2020) using a 1 Mb window. We also annotated other publicly available alfalfa genomes including XJDY (Chen et al. 2020), Zm1 (Shen et al. 2020), and Zm4 (Long et al. 2022) for R genes using the same pipeline. After identifying and annotating R genes, we examined the number of clusters formed across different domain classes. An R-gene cluster was defined within each genome, haplotype, chromosome, and NLR subclass (e.g., TNL, CNL, RNL, and RxNL) as a run of three or more R genes with adjacent genes located ≤ 200 kb apart, measured between gene midpoints.

### Comparative genomic analyses

We used python JCVI-MCScanX (Python version; Tang et al. 2024) to identify gene-based synteny between the four haplotypes of Regen-SY27x using default parameters (Supplemental Methods). The Regen-SY27x genome was also compared with the first haplotype of XJDY (Chen et al. 2020) using the same pipeline. We used OrthoFinder v3.1.0 (Emms and Kelly 2019) to identify gene orthogroups among the alfalfa genomes; Regen-SY27x, Zm1 (Shen et al. 2020), Zm4 (Long et al. 2022), XJDY (Chen et al. 2020); and five other legume species, *M. ruthenica* (Wang et al. 2021), *M. truncatula* v5.0 (Pecrix et al. 2018), *Pisum sativum* v1a (Kreplak et al. 2019)*, Cicer arietinum* cv. CDC Frontier v1.0 (Varshney et al. 2013), and *Trifolium pratense* (De Vega et al. 2015) using the annotated primary protein sequences. The orthologous gene families and phylogenetic tree topology inferred from OrthoFinder v3.1.0 (Emms and Kelly 2019) were input to CAFE5 v5.1.0 (Mendes et al. 2021) to identify significant expansion or contraction in each gene family (P < 0.01). We used Mummer v4.0.0beta2 (Marçais et al. 2018) to sequentially align the four haplotypes and its output was used in Synteny and Rearrangement Identifier (SyRI) v1.7.1 (Goel et al. 2019) to profile structural variation (SV) between the pseudo-chromosomes of the four haplotypes. Other available high-quality, allele aware and chromosome-scale alfalfa genomes including XJDY (Chen et al. 2020), ZM1 (Shen et al. 2020), and Zm4 (Long et al. 2022) were also compared with the Regen-SY27x genome assembly using minimap2 v2.26 (Li 2018) and SyRI v1.7.1 (Goel et al. 2019) to profile sequence and structural variants between them (see Supplementary Methods). The identified SVs were visualized using plotsr v1.1.5 (Goel and Schneeberger 2022).

### Data visualization and genome browser

To visualize the data and examine assembly collinearity compared to the other alfalfa genomes in the Legume Information System Datastore (https://data.legumeinfo.org/), dotplots were created using DotPlotly. The Regen-SY27x assembly and the final structural annotations were added as a set of per-haplotype genomes to a JBrowse2 (Diesh et al. 2023) instance running at the Legume Information System (https://medicago.legumeinfo.org/tools/jbrowse2/?session=share-zp6qTP7cv_&password=kAA1r).

## Results and Discussion

### Assembly contiguity and length

The reference genome for Regen-SY27x was assembled using 92,918,416,171 bp of sequences from circular consensus PacBio HiFi reads after quality filtering, providing ∼116× haploid genome coverage with an average read length of 15,989 bases (Supplementary Table 1 and Supplementary Table 2). For scaffolding, we generated 94,338,638 Omni-C read pairs (2 x 150 bp), providing a total of 28.3 Gb of data. Further haplotype phasing and pseudochromosome-level scaffolding were performed using alleles phased in the linkage groups (Supplementary Table 3) with the occurrence of those same alleles in the genome assembly (see details in the Material and Methods section). The Regen-SY27x genome was assembled into four phased sets. We defined the total haplotype for each set as the sum of all phased sequences (pseudochromosomes plus unplaced scaffolds), which ranged in size from 789.2 Mb (haplotype 1) to 914.3 Mb (haplotype 4). From these, we identified the pseudochromosome assembly, consisting of the eight scaffolds per haplotype. These 32 pseudochromosomes represent the anchored portion of the genome, totaling 2.59 Gb with a scaffold N50 of 83.3 Mb. The final assembly contained four haplotypes for each chromosome. Assembly statistics including both the full assembly (pseudochromosomes + unplaced scaffolds) as well as the pseudochromosomes are shown in Table 1.

**Table 1.** Assembly statistics per haplotype for the full assembly and the pseudochromosome assembly of Regen-SY27x.

| Group | Metric | Total<br>haplotypes | Pseudo-<br>haplotype 1 | Pseudo-<br>haplotype 2 | Pseudo-<br>haplotype 3 | Pseudo-<br>haplotype 4 |
| --- | --- | --- | --- | --- | --- | --- |
| Pseudochromosomes + unplaced scaffolds | Scaffolds | 6,021 | 1,043 | 1,033 | 1,035 | 2,910 |
|  | Contigs | 7,162 | 1,331 | 1,348 | 1,306 | 3,177 |
|  | Scaffold Length | 3,286,073,882 | 789,244,119 | 789,460,494 | 793,088,424 | 914,280,845 |
|  | Contig Length | 3,285,806,182 | 789,179,319 | 789,386,594 | 793,019,724 | 914,220,545 |
|  | Contig N50 | 3,406,605 | 4,276,997 | 2,935,843 | 3,505,361 | 3,106,875 |
|  | Contig L50 | 254 | 55 | 68 | 58 | 76 |
|  | Scaffold N50 | 79,699,470 | 79,699,470 | 73,507,436 | 75,653,446 | 81,593,309 |
|  | Scaffold L50 | 19 | 5 | 5 | 5 | 6 |
|  | Max scaffold | 96,423,042 | 96,190,425 | 90,827,970 | 91,408,138 | 96,423,042 |
|  | Min scaffold | 10,665 | 11,938 | 11,779 | 11,494 | 10,665 |
| Pseudochromosomes | Scaffolds | 32 | 8 | 8 | 8 | 8 |
|  | Contigs | 1,141 | 284 | 314 | 272 | 271 |
|  | Scaffold Length | 2,594,518,724 | 653,415,314 | 625,696,371 | 648,084,291 | 667,322,748 |
|  | Contig Length | 2,594,267,024 | 653,356,514 | 625,626,971 | 648,019,091 | 667,264,448 |
|  | Contig N50 | 4,725,360 | 4,956,865 | 4,321,830 | 4,725,360 | 4,642,456 |
|  | Contig L50 | 170 | 41 | 46 | 41 | 44 |
|  | Scaffold N50 | 83,291,558 | 83,716,887 | 83,291,558 | 86,231,604 | 83,189,900 |
|  | Scaffold L50 | 15 | 4 | 4 | 4 | 4 |
|  | Max scaffold | 96,423,042 | 96,190,425 | 90,827,970 | 91,408,138 | 96,423,042 |
|  | Min scaffold | 62,554,310 | 69,581,426 | 62,554,310 | 69,463,442 | 71,615,600 |

The total assembly length was 3.2 Gb, consistent with the expected ∼800 Mb genome size per haplotype (Table 1 and Supplementary Table 4). Assembly lengths and contiguity were generally balanced across the four haplotype pseudochromosomes. However, inclusion of unplaced scaffolds makes haplotype 4 approximately 15% larger, with more than twice the number of contigs and nearly three times the number of scaffolds compared to the next largest haplotype, although N50 values remain comparable. This pattern likely reflects the original hifiasm partitioning, which generated many small scaffolds in haplotype 4. Because scaffolds that could not be assigned to a haplotype based on the genetic map were retained in their original haplotype, haplotype 4 carries a disproportionate number of small unplaced scaffolds, some of which may belong to other haplotypes.

Overall assembly contiguity of the genome is high. Scaffold N50s are close to chromosome lengths due to pseudomolecule construction (Table 1). Contig N50s, which better reflect the segmentation of the genome, range from 2.9 to 4.3 Mb in individual haplotypes and 3.4 Mb when all haplotypes are combined when considering pseudochromosomes + unplaced scaffolds. Contig N50s for only pseudochromosomes range from 4.3 to 5.0 Mb. The larger contig N50s in the pseudochromosomes compared to the pseudochromosomes + unplaced scaffolds reflects the difficulty in placing the smaller scaffolds using the genetic map. In short, the assembly statistics show that the assembly is close to the expected genome size, is highly contiguous, and is well partitioned among haplotypes. Across the 32 pseudochromosome-scale scaffolds (representing four phased haplotypes for Chromosomes 1–8), GC content calculated in 1 Mb windows ranges from 23.7% to 50.4%, with a mean of 34.2%. It is largely uniform along chromosome arms with localized depressions that coincide with repeat-rich blocks (Fig. 1).

**Figure 1.**
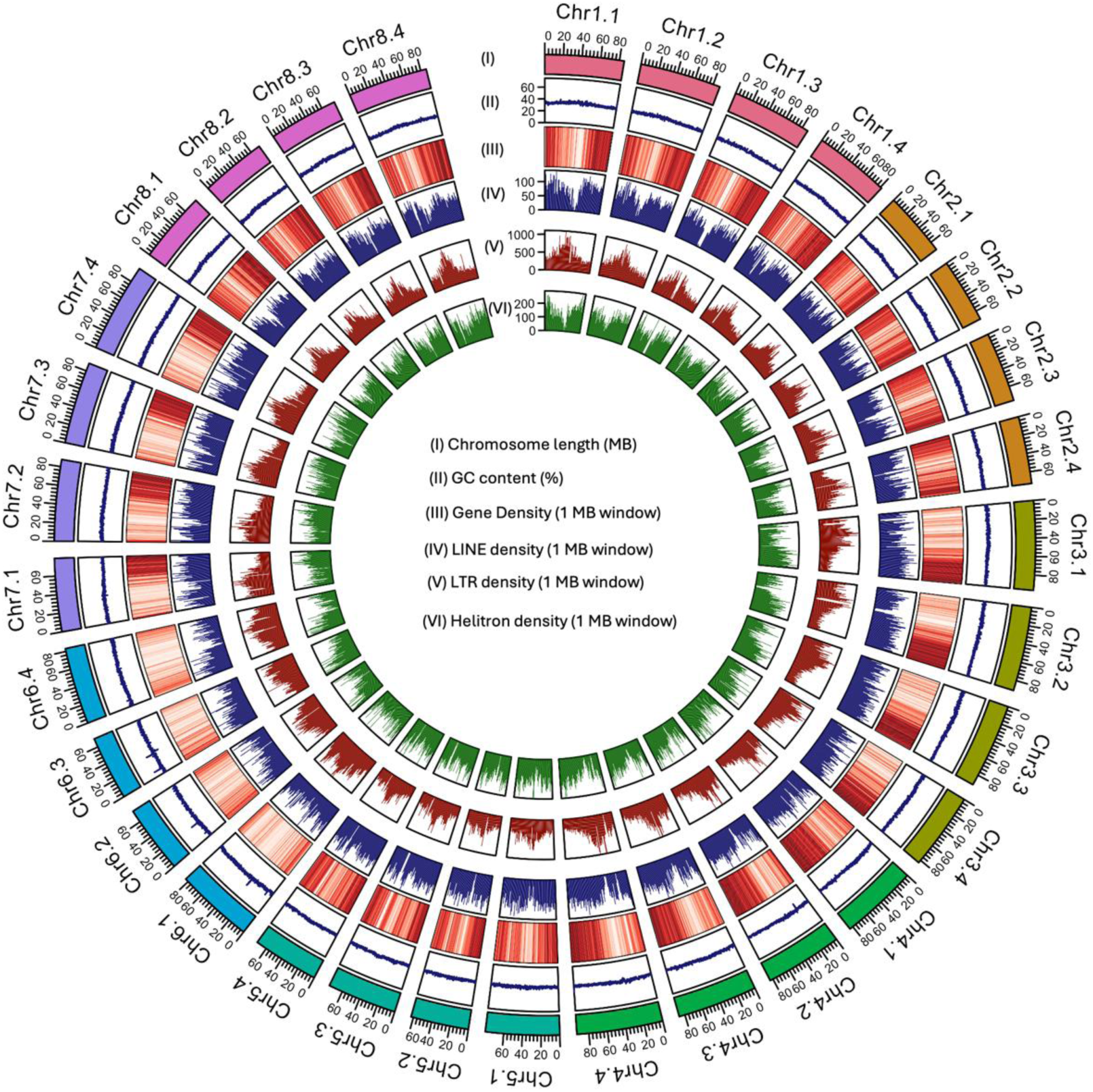
Circos plot of the phased Regen-SY27x genome. Tracks from outermost to innermost: (I) chromosome lengths, (II) % GC content, (III) gene density heatmap per 1 MB block, darker red color corresponds to higher gene density, (IV) LINE density barplots per 1 MB block, (V) LTR density barplots per 1 MB block, and (VI) Helitron content barplots per 1 MB block.

The Regen-SY27x assembly represents a sophisticated approach to autotetraploid genome reconstruction, leveraging both physical proximity data and genetic inheritance patterns to create a highly accurate and fully phased chromosome scale assembly. To maximize physical contiguity, we utilized Omni-C^TM^ instead of traditional restriction-enzyme-based Hi-C. The sequence-independent nature of DNase I digestion in the Omni-C protocol provided more uniform coverage across the repetitive alfalfa landscape and eliminates the blind spots associated with fixed restriction sites (Ramani et al. 2016; Ma et al. 2018). This ensured a high density of physical anchors in both gene-rich distal arms and repetitive pericentromeric regions (Gridina et al. 2021). Our novel assembly strategy integrated Omni-C^TM^ data for long-range physical scaffolding with fully phased SNP-based, dosage-aware linkage maps to achieve precise phasing of the Regen-SY27x haplotypes. This dual-evidence approach mirrors recent successes in other complex polyploids, such as hexaploid sweet potato, where simplex markers from linkage map and Hi-C data were integrated to overcome the limitations of traditional scaffolding (Wu et al. 2025). This approach prevents common assembly pitfalls like haplotype collapse and ensures both structural integrity and biological accuracy by integrating physical and recombination links. Previous alfalfa genome assemblies have used linkage maps primarily for validation of assembly accuracy, rather than to guide haplotype phasing or to detect and resolve chimeric scaffolds during the assembly process (Chen et al. 2020; Shen et al 2020).

### Haplotype partitioning assessment using kmer plots

The kmer comparison plots, which compare the kmer content in the reads with that of the assembly, are important genome assembly assessment tools. These graphs provide coverage information which allow for the disambiguation of real kmers and those which arise from sequencing errors (Mapleson et al. 2017). In addition, the kmer peaks indicate the ploidy of the genome, reflect haplotype similarity, and allow for the confirmation of copy number calls in the assembly (Carvalho et al. 2016).

The kmer spectra comparison (Fig. 2) confirmed the high completeness and precise partitioning of the Regen-SY27x assembly. In the full assembly plot (Fig. 2A), the clear separation between the error peak (low-frequency black kmers) and the alfalfa genomic peaks confirms sufficient coverage for high-fidelity assembly. The assembly effectively excludes sequencing errors while capturing true genomic kmers in their expected multiplicities. Notably, the 1X (red) peak is the most prominent, containing significantly more kmers than the shared 2X, 3X, or 4X peaks. This indicates that a large proportion of the alfalfa genome is highly polymorphic, with unique kmers distinguishing the four haplotypes despite their autotetraploid origin.

**Figure 2.**
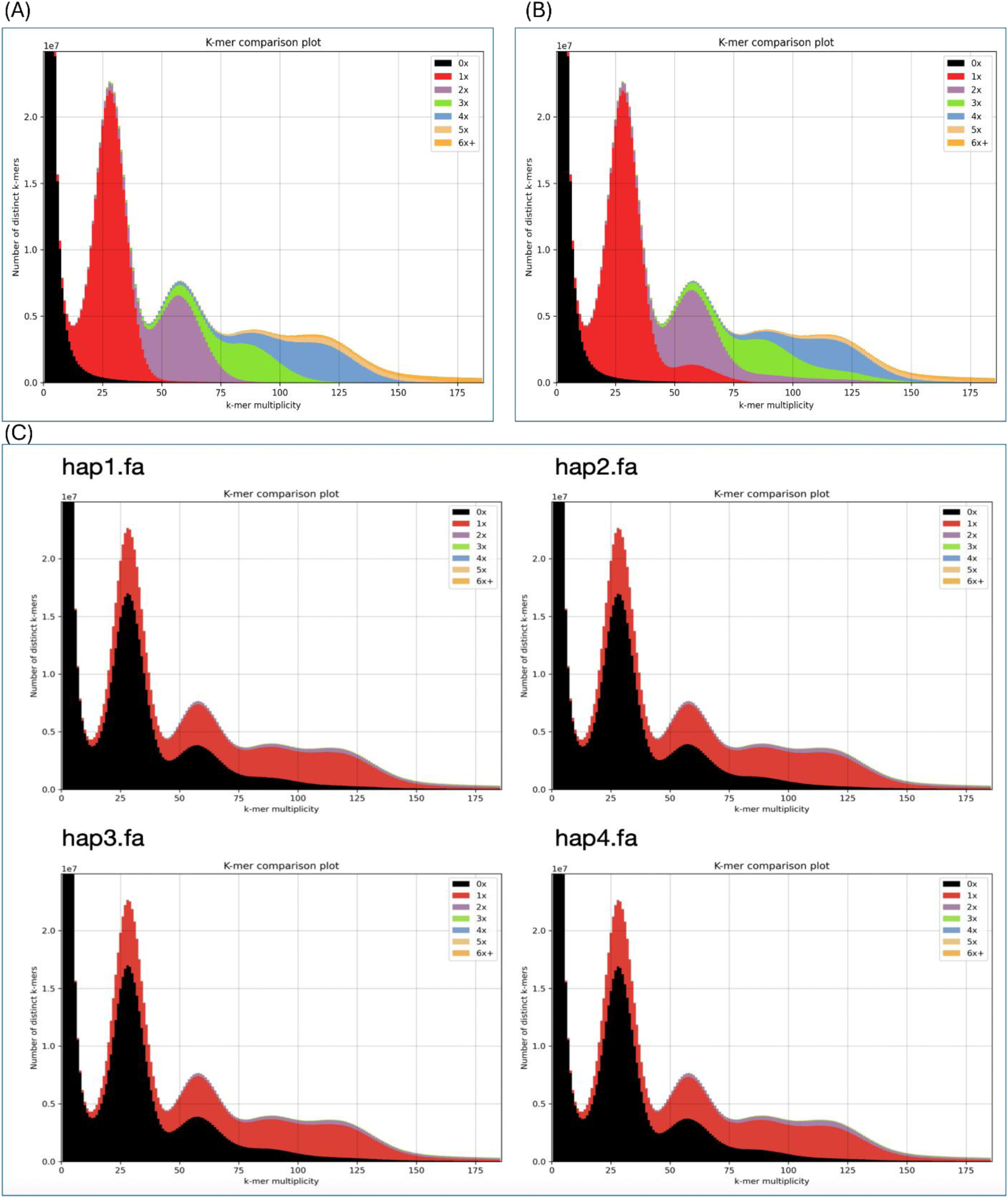
Kmer plots, including both pseudochromosomes and unplaced scaffolds, generated by the Kmer Analysis Toolkit (KAT) for A) the phased Regen-SY27x assembly containing all four haplotypes, B) the phased Regen-SY27x assembly containing all four haplotypes before the final phasing parameters in hifiasm were achieved, C) kmer plots of individual haplotypes.

Individual haplotype plots (Fig. 2C) further validate the partitioning by demonstrating that each assembly partition captures the expected ∼25% of the unique genomic kmers. This distribution, representing the expected haplotypic proportions, confirms that we successfully resolved the four homologs. Compared to earlier assembly iterations (Fig. 2B), these results demonstrate that the chosen parameters in hifiasm and use of linkage maps successfully minimized haplotype collapse and maximized structural resolution.

### Chimeric scaffolds breakdown using dotplots

The HiFi-based primary contig assemblies achieved a level of contiguity that formerly required iterative scaffolding, and Omni-C proximity ligation and phased linkage maps enabled us to develop phased chromosomal pseudomolecules. Nevertheless, the occurrence of chimeric sequences underscores the necessity of rigorous manual curation (Jiao et al. 2017; Sato et al. 2021; Marcolungo et al. 2023). By utilizing high-quality reference genomes or established chromosomal assemblies, the mis-joins between haplotypes were identified and corrected to ensure structural accuracy. We utilized the *M. sativa* subsp. *caerulea* PI 464715 genome assembly (Li et al. 2020) to assist in Regen-SY27x assembly and phasing. PI 464715 is recognized as a diploid progenitor of autotetraploid alfalfa (Small and Jomphe 1989). The genome assembly of PI 464715 was constructed using Oxford Nanopore Technology, yielding a high-quality reference where 98.5% of the contigs are anchored across eight pseudochromosomes with a BUSCO completeness of 97.7%. Dotplots showing alignments of scaffolds contributing to pseudochromosomes for each haplotype to the PI464715 assembly were used to visually assess the completeness and any redundancy of each individual haplotype, as well as to identify potential breakpoints for scaffolds that are indicated as chimeric by the genetic map (Fig. 3). The dotplots show that these scaffolds and the resulting pseudochromosome cover most of the PI464715 genome with little redundancy. Scaffolds were placed into the pseudomolecule by the order and orientation suggested by the dot plot. Structural variation in Regen-SY27x compared to PI464715, with the exception of chimeras identified by the genetic-map, were not altered as they are likely true structural differences.

**Figure 3.**
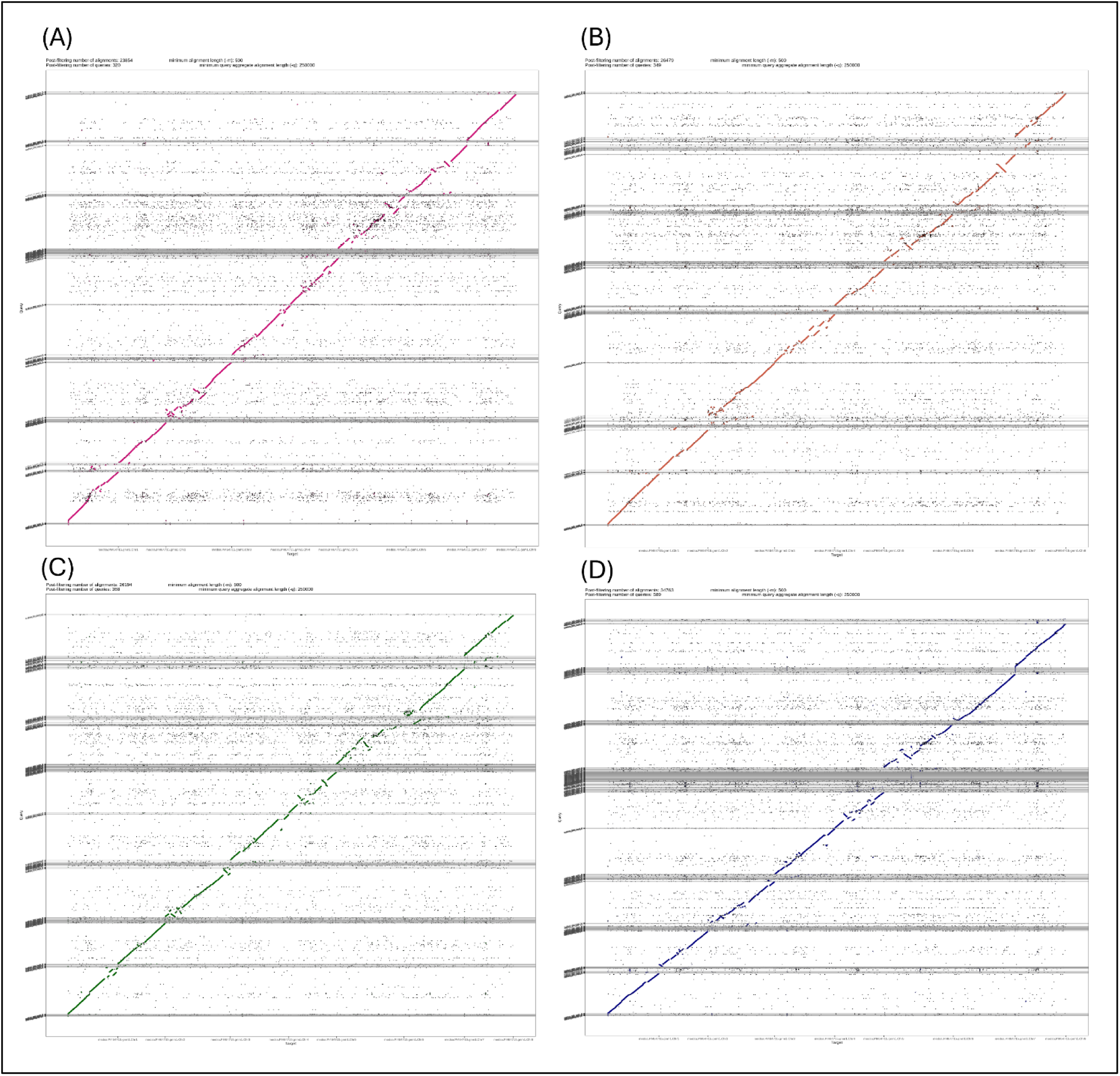
Dotplots of each of the haplotypes from the final assembly (y axis) against PI464715 (x axis). A) Haplotype 1, B) Haplotype 2, C) Haplotype 3, D) Haplotype 4.

### Annotation

After removing redundancy with our adjudication method, there were a total of 221,688 genes in the assembly, including 54,412 genes, 55,178 genes, 54,512 genes, and 57,586 genes in haplotypes 1 to 4, respectively. More than 99% of genes in each haplotype were incorporated into the haplotype’s pseudochromosomes, leaving less than 1% of genes for each haplotype in the unplaced scaffolds.

Repeat annotation of the Regen-SY27x genome revealed that 62.65% of the assembly consists of interspersed repeats, dominated by long terminal repeat (LTR) retrotransposon (36.7%) (Table 2). Among LTRs, Gypsy (13.97%), Copia (8.36%), and a large unclassified fraction (14.29%) represented the most abundant categories. Helitrons occupied 5.52% of the assembly, highlighting their high abundance in alfalfa. In comparison, average Helitron content across 26 Fabaceae genomes is 1.38% (Li et al. 2023), indicating that alfalfa harbors a relatively high proportion. Helitrons are unique among DNA transposons in their ability to capture, duplicate, and shuffle host gene fragments, often creating chimeric structures or novel paralogs (Kapitonov and Jurka 2007). In plants, such activity has been directly linked to gene family diversification. For example, in maize, Helitrons have duplicated fragments of disease-resistance genes and other genic regions, contributing to haplotype variation and lineage-specific expansions (Lai et al. 2005; Morgante et al. 2005; Yang and Bennetzen 2009).

**Table 2:** Repeat classes summary in the Regen-SY27x genome.

| Class | Sub-class | Count | Size (bp) | %Genome |
| --- | --- | --- | --- | --- |
| DNA Transposon |  | 71,396 | 14,139,653 | 0.43 |
| LINE | CR1 | 43 | 6,834 | 0.00 |
|  | CRE | 1,551 | 644,046 | 0.02 |
|  | Jockey | 78 | 15,869 | 0.00 |
|  | L1 | 210,534 | 101,514,852 | 3.09 |
|  | L2 | 581 | 180,250 | 0.01 |
|  | RTE | 48,040 | 11,258,729 | 0.34 |
| LTR | Bel Pao | 1,254 | 603,559 | 0.02 |
|  | Copia | 310,338 | 275,092,005 | 8.37 |
|  | Endogenous Retrovirus | 2,033 | 337,365 | 0.01 |
|  | Gypsy | 466,960 | 458,992,456 | 13.97 |
|  | Unknown | 570,137 | 469,594,319 | 14.29 |
| SINE | ID | 30,672 | 3,066,228 | 0.09 |
|  | tRNA | 44,974 | 13,370,911 | 0.41 |
|  | Unknown | 722 | 103,917 | 0.00 |
| TIR | CACTA | 48,657 | 18,355,896 | 0.56 |
|  | KDZ | 195 | 57,049 | 0.00 |
|  | Kolobok | 27 | 3,483 | 0.00 |
|  | Mutator | 287,658 | 109,722,958 | 3.34 |
|  | PIF Harbinger | 123,440 | 33,826,478 | 1.03 |
|  | PILE | 13 | 2,110 | 0.00 |
|  | Sola | 146 | 70,936 | 0.00 |
|  | Tc1 Mariner | 85,947 | 21,953,850 | 0.67 |
|  | Transib | 27 | 10,649 | 0.00 |
|  | hAT | 137,798 | 40,230,169 | 1.22 |
|  | piggyBac | 2,903 | 337,892 | 0.01 |
|  | polinton | 1,748 | 267,494 | 0.01 |
| non-LTR | Ngaro_YR | 5,022 | 1,160,533 | 0.04 |
|  | pararetrovirus | 8,577 | 4,901,570 | 0.15 |
| non-TIR | Crypton_YR_transposon | 203 | 72,767 | 0.00 |
|  | Helitron | 535,577 | 181,232,884 | 5.52 |
| rDNA 45S |  | 6,541 | 27,284,625 | 0.83 |
| Repeat Fragment |  | 1,034,798 | 264,973,241 | 8.06 |
| snDNA |  | 8,357 | 4,703,100 | 0.14 |
| snRNA |  | 1,793 | 353,874 | 0.01 |
| Total |  | 4,048,740 | 2,058,442,551 | 62.65 |

Abbreviations: LINE, Long Interspersed Nuclear Element; L1, LINE-1 family; RTE, Retrotransposable Element; LTR, Long Terminal Repeat retrotransposon; SINE, Short Interspersed Nuclear Element; TIR, Terminal Inverted Repeat transposon; CACTA, CACTA (En/Spm) transposon; PILE, Plant Inverted-repeat Low-copy Element (PIF-like); POLE, Pong-like element (Harbinger-related); snDNA, satellite DNA.

The TEs divergence landscape revealed a sharp low-divergence peak (<2%) for LTRs, indicating a recent burst of retrotransposon activity, superimposed on a broader 10 to 20% shoulder consistent with earlier amplification waves (Supplementary Fig. 1). These results are consistent with earlier alfalfa genome assemblies (Chen et al. 2020; Shen et al. 2020). Helitrons and TIR superfamilies spanned wide divergence bins, suggesting long-running, lower-level mobilization.

Gene density (Fig. 1, track 3) peaks on distal chromosome arms and is consistently reduced in the broad central regions, which also exhibit elevated LTR signals (Fig. 1, track 4), indicating putative peri-/centromeric regions. In contrast, Helitron patterns (Fig. 1, track 6) show an opposite trend relative to LTRs, suggesting their potential enrichment in or near genic regions. These spatial patterns are highly concordant among the four haplotypes for each chromosome, supporting successful phasing and a structurally consistent assembly without large-scale anomalies or GC spikes indicative of contamination.

### Completeness and copy number of universal orthologs

The BUSCO was run on both the genome assembly and the annotated genes and, like other technical validations, was used iteratively to improve the assembly and annotation. Both the genome and annotated genes indicated that nearly all the universal ortholog genes captured in the genome assembly were annotated. Fabales Odb10 dataset in BUSCO was used to search for universal orthologs in the Fabales. The number of captured BUSCOs in the full assembly (99.3%) and each individual haplotype (haplotype1 94.9%, haplotype2 95.9%, haplotype3 94.4%, haplotype4 96.9%) shows that the full assembly and each of the haplotypes has a high degree of completeness. As expected in a phased assembly with multiple haplotypes, nearly all (98.6%) of the BUSCOs were duplicated within the assembly. These results are consistent with other haplotype-resolved autotetraploid assemblies of alfalfa (Chen et al. 2020; Long et al. 2022) and potato (Bao et al. 2022; Hoopes et al. 2022; Sun et al. 2022), which show 90 to 97.4% duplicated genes.

Approximately 75% of the found BUSCOs were present in exactly four copies. In addition, 82% of found BUSCOs had at least one copy per haplotype. These results offer a compelling contrast to other autotetraploid models, such as the potato genome. The haplotype-resolved potato genome assemblies (Sun et al. 2022; Godec et al. 2025) reported that only 53.6-55.3% of genes were present in all four haplotypes. This suggests that while potato may undergo more rapid or extensive hemizygous gene loss or structural divergence, the alfalfa genome retains a bigger core gene set across its four homologous chromosomes. Taken together, the BUSCO results suggest that the full assembly, as well as each of the individual haplotypes, is highly complete.

### Nucleotide-binding leucine-rich repeats (NLR) gene annotation

Nucleotide-binding leucine-rich repeat (NLR) genes form a major superfamily of disease resistance (R) genes and are defined by the presence of the conserved NB-ARC domain at their core, which functions as the molecular switch for signaling (Fig. 4). NLRs are further classified according to their N-terminal domains into coiled-coil like (CC–NB-ARC–LRR/CNLs), Toll/Interleukin-1 receptor like (TIR–NB-ARC–LRR/TNLs), RPW8-like (RPW8–NB-ARC–LRR/RNLs), and Rx-type (Rx–NB-ARC–LRR/RxNLs), all of which represent complete proteins containing an N-terminal signaling domain, the central NB-ARC domain, and a C-terminal LRR region (Shao et al. 2016). In contrast, truncated or partial NLRs lack one or more of these domains and are subdivided into CNB (CC+NB-ARC only), RxNB (Rx+NB-ARC only), TNB (TIR+NB-ARC only), and RNB (RPW8+NB-ARC only). Additionally, some sequences were classified as NB-ARC only when the central domain was present without clearly associated N- or C-terminal extensions. This domain-based classification highlights the conserved role of the NB-ARC as the core signaling hub of all NLRs, while also capturing the structural diversity and evolutionary flexibility of the superfamily (Meyers et al. 2003; Jones et al. 2016; Tamborski and Krasileva 2020). We used this classification to annotate the NLR genes in alfalfa genomes using *de novo* FindPlantNLRs (Chen et al. 2022) pipeline. Additionally, NLRs were classified in three super categories based on the presence or absence of canonical domains at the N-terminus, central NB-ARC, and C-terminal LRR regions. Genes containing all three major domains were categorized as complete NLRs, further subdivided into CNL, TNL, RNL, and RxNL. Genes missing one or more of the canonical domains were placed into partial classes, including CNB, RxNB, RNB, and TNB. Genes with only the conserved NB-ARC domain, or NB-ARC with partial NLR signatures, were grouped as the unknown class. This hierarchical classification enabled us to capture both the canonical full-length NLRs and truncated or atypical variants, providing a comprehensive overview of R-gene diversity in autotetraploid alfalfa.

**Figure 4.**
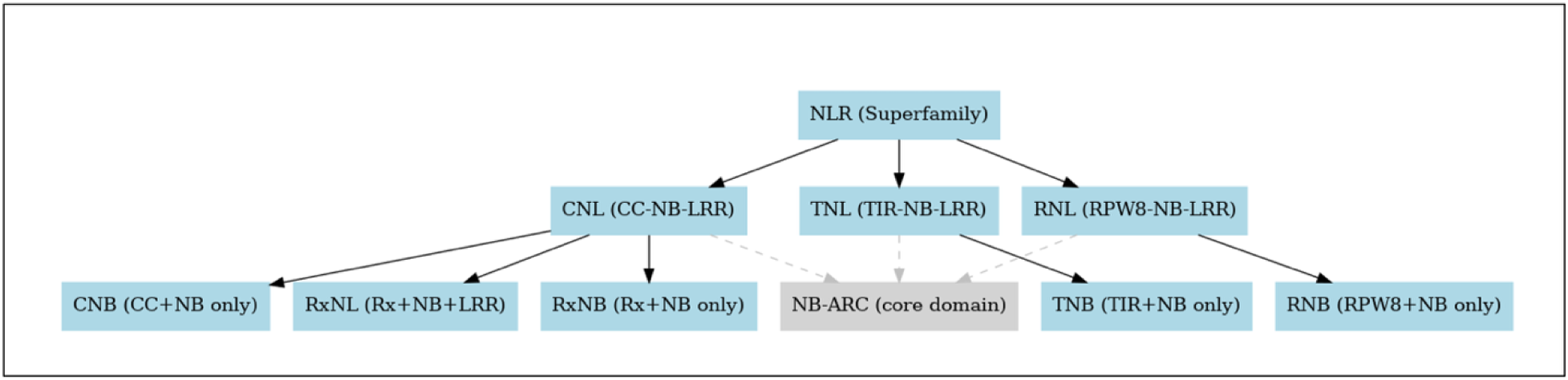
Nucleotide binding leucine-rich repeats (NB-LLR/ NLR) gene superfamily classification.

A total of 3,696 NLR genes were identified in the Regen-SY27x assembly, distributed across four haplotypes (Supplementary Table 5). Among these, 2,513 genes belonged to the complete classes (CNL, RxNL, RNL, and TNL), while 714 were assigned to the unknown class (NB-ARC/NB-ARC+NLR only) and 469 to partial classes. Within the complete categories, TNL (1,032) and RxNL (1,024) represented the largest groups, followed by CNL (424), whereas RNL genes (33) were the least abundant. Partial classes were comparatively smaller, including CNB (49), RxNB (164), RNB (6), and TNB (250) (Table 3).

**Table 3:** The resistance (R) gene domain clusters identified in the genome of Regen-SY27x using the FindPlantNLRs pipeline.

| Class | No. of<br>R genes | No. of<br>clusters | % of R<br>genes in<br>clusters | Largest<br>cluster | No. of<br>R genes | No. of<br>clusters | % of R<br>genes in<br>clusters | Largest<br>cluster | No. of<br>R genes | No. of<br>clusters | % of R<br>genes in<br>clusters | Largest<br>cluster | No. of<br>R genes | No. of<br>clusters | % of R<br>genes in<br>clusters | Largest<br>cluster |
| --- | --- | --- | --- | --- | --- | --- | --- | --- | --- | --- | --- | --- | --- | --- | --- | --- |
|  | Haplogroup 1 |  |  |  | Haplogroup 2 |  |  |  | Haplogroup 3 |  |  |  | Haplogroup 4 |  |  |  |
| CNL | 109 | 6 | 50 | 17 | 112 | 7 | 54 | 17 | 98 | 5 | 42 | 15 | 105 | 6 | 45 | 15 |
| RNL | 8 | 1 | 38 | 3 | 11 | 1 | 82 | 3 | 8 | 1 | 38 | 3 | 6 | 1 | 50 | 3 |
| RxNL | 279 | 18 | 63 | 24 | 242 | 11 | 62 | 14 | 256 | 17 | 59 | 12 | 247 | 15 | 60 | 12 |
| TNL | 264 | 14 | 70 | 25 | 271 | 15 | 70 | 17 | 234 | 10 | 73 | 18 | 263 | 13 | 66 | 24 |
| CNB | 12 | 0 | 0 | 0 | 11 | 0 | 0 | 0 | 10 | 0 | 0 | 0 | 16 | 0 | 0 | 0 |
| RNB | 3 | 0 | 0 | 0 | 1 | 0 | 0 | 0 | 1 | 0 | 0 | 0 | 1 | 0 | 0 | 0 |
| RxNB | 52 | 3 | 29 | 8 | 33 | 0 | 0 | 0 | 42 | 1 | 21 | 6 | 37 | 1 | 11 | 4 |
| TNB | 53 | 2 | 28 | 5 | 62 | 5 | 34 | 4 | 66 | 3 | 53 | 8 | 69 | 4 | 28 | 5 |
| NBARC/NLR | 190 | 11 | 34 | 9 | 181 | 13 | 42 | 7 | 173 | 8 | 35 | 8 | 170 | 8 | 34 | 8 |

Abbreviations: CNB = Coiled-coil Nucleotide-Binding, CNL = Coiled-coil Nucleotide-Binding Leucine-rich repeat, NBARC = Nucleotide-Binding domain shared among APAF-1, R gene products, and CED-4, NLR = Nucleotide-binding site Leucine-rich Repeat, RNB = Resistance Nucleotide-Binding, RNL = Resistance Nucleotide-Binding Leucine-rich repeat, RxNB = Rx-type Nucleotide-Binding, RxNL = Rx-type Nucleotide-Binding Leucine-rich repeat, TNB = Toll/interleukin-1 Receptor Nucleotide-Binding, and TNL = Toll/interleukin-1 Receptor Nucleotide-Binding Leucine-rich repeat.

The distribution of NLR classes showed both haplotype-level similarities and genotype specific differences (Fig. 5, Table 3). The highest number of R genes was identified in the Regen-SY27x genome followed by XJDY, Zm4, and Zm1 (Supplementary Table 5). The relative proportions of the major complete NLR subclasses were broadly conserved, with RxNL and TNL consistently representing the dominant fractions across all haplotypes and genotypes.

**Figure 5.**
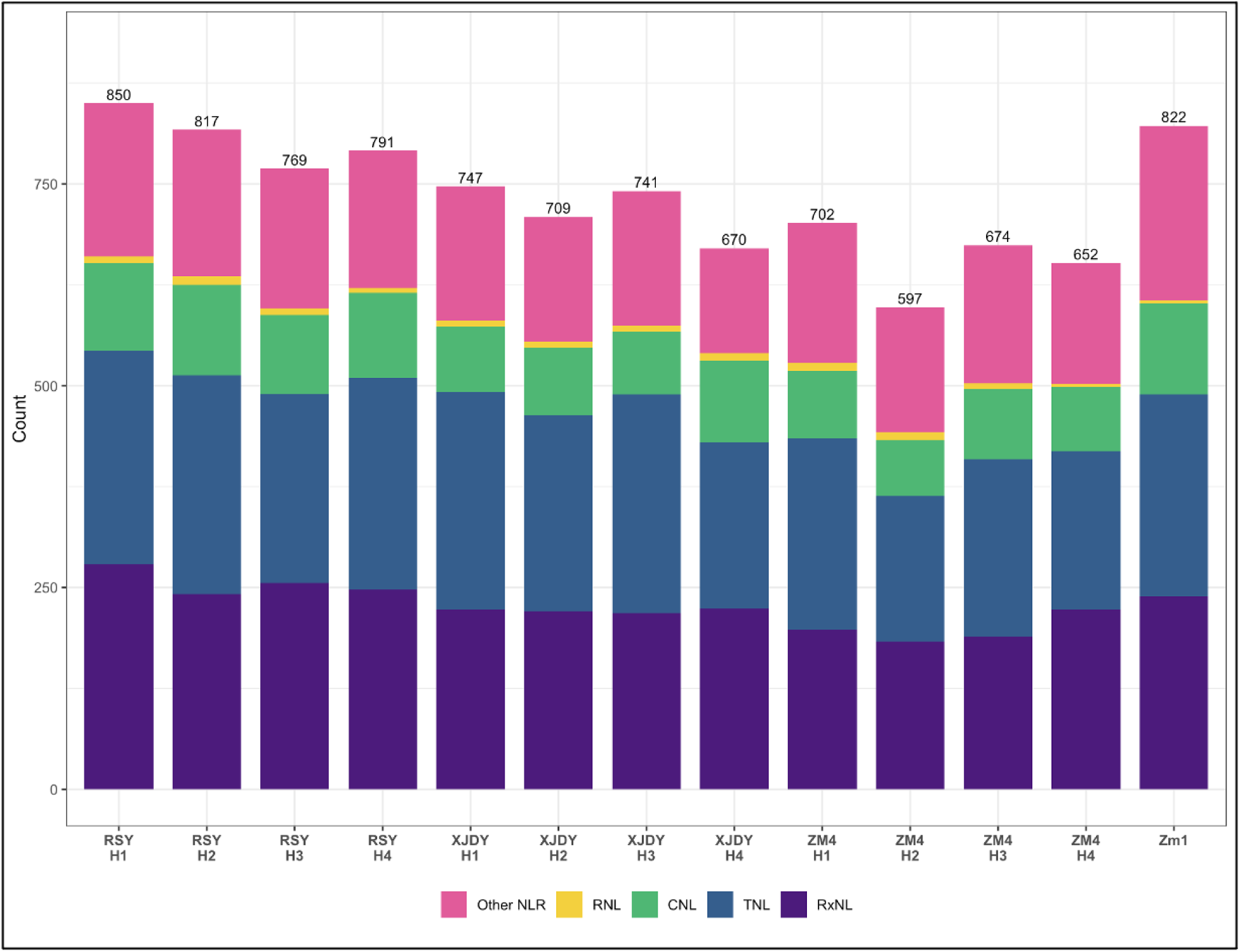
Number of nucleotide binding and leucine-rich repeat (NLR) genes recovered from four alfalfa genomes (Regen-SY27x, RSY; XJDY, Zm4, and Zm1) in four specific categories, RNL, CNL, TNL, and RxNL, plus unclassified categories.

Clustering analysis revealed that a substantial proportion of NLRs occur in tandemly arranged clusters across all four Regen-SY27x haplotypes (Table 3). The largest clusters were found on distal arms of chromosome 3, 4, 6, and 8 (Fig. 6). Among the complete classes, TNLs and RxNLs were the most frequently clustered, with ∼60 to 73% of genes found in clusters and the largest clusters containing up to 25 members. CNLs also showed moderate clustering, with ∼42 to 54% of genes arranged in clusters and maximum cluster sizes of 15 to 17 genes. In contrast, RNLs were rare but showed variable clustering. Across all four alfalfa autotetraploid genomes analyzed (Regen-SY27x, XJDY, Zm4, and Zm1), we found that TNLs and RxNLs consistently showed the strongest clustering (∼60 to 75%), often forming large tandem arrays, whereas CNLs exhibited intermediate clustering (∼37 to 50%) and RNLs were rare and variably clustered (Supplementary Tables 6-8). This conserved subclass-specific clustering pattern suggests that tandem duplication is a predominant driver of R-gene diversification in autotetraploid alfalfa. These results are in line with previous work in diploid relatives. In *M. truncatula*, Ameline-Torregrosa et al. (2008) identified 333 non-redundant NBS-LRR genes, with TNLs comprising ∼70% of the repertoire and showing frequent tandem organization. Likewise, in *M. ruthenica*, a recent genome-wide study reported 338 NLRs dominated by TNL and CNL subclasses, again enriched in clusters (Tong et al. 2022). Together, these comparisons indicate that a TNL-rich, cluster-prone NLR landscape is an evolutionarily conserved hallmark of *Medicago*, extending from diploid species to autotetraploid cultivars. Our findings further suggest that the expansion of TNL and RxNL clusters in alfalfa has been maintained despite polyploidization, underscoring their importance as dynamic reservoirs of resistance diversity.

**Figure 6.**
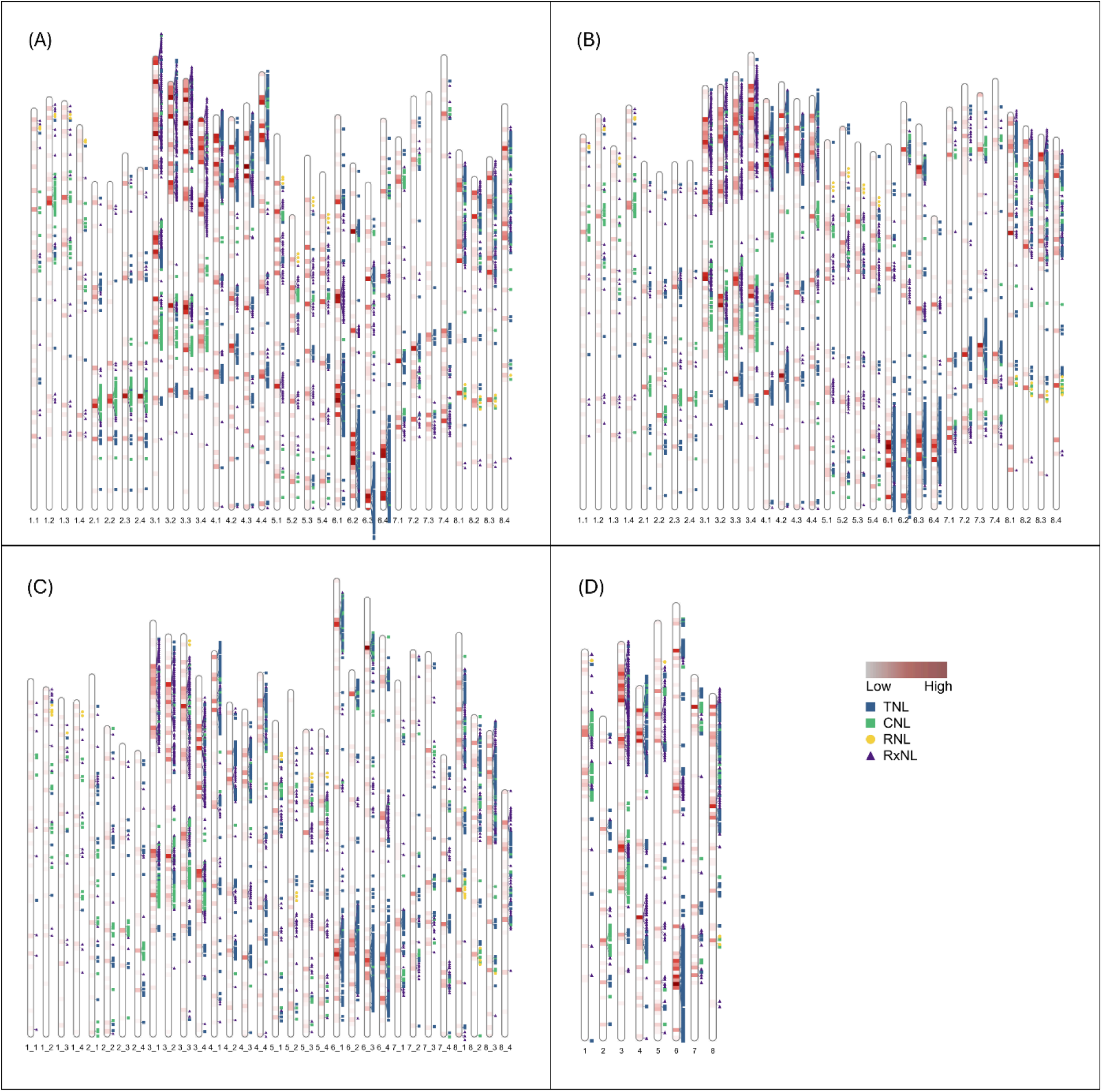
Locations of R genes on the chromosomes of (A) Regen-SY27x, (B) XinJiang DaYe, (C) ZhongmuNo.4, and (D) ZhongmuNo.1 genomes. Density of R genes is shown as a heat map in 1 Mb segments in chromosome karyotypes, with lighter red color indicating fewer genes and darker color indicating more genes. The blue box, green box, yellow circle, and purple triangle alongside chromosome karyotypes display positions of N-terminal domains of **CNL** - Coiled-coil NLR (CC–NB-ARC–LRR), **TNL** - TIR-domain NLR (TIR–NB-ARC–LRR; TIR = Toll/Interleukin-1 receptor homology), **RNL**-RPW8-like NLR (RPW8-like CC–NB-ARC–LRR; “helper” NLRs), and **RxNL**-Rx-type N-terminal domain + NB-ARC–LRR (complete Rx-NLR) genes.

### Comparative genomic analyses

Across the four phased haplotypes of Regen-SY27x, gene-based macrosynteny evaluated using MCScan was strongly conserved chromosome-by-chromosome (Fig. 7). Across all pairwise haplotype comparisons (Supplementary Figs. 2A, 2B and 2C), the majority of genes exhibit a 1:1 syntenic depth (82 to 83%), indicating strong single-copy conservation between haplotypes, with only a minor fraction showing absence of synteny or duplication, which is consistent with BUSCO analyses results. Within Regen-SY27x, haplotype comparisons (Fig. 7) show many (>665) syntenic blocks with a low median gene number (9 to 10) per block indicative of fragmented collinearity. Macrosynteny of haplotypes 1 of Regen-SY27x and XJDY (Supplementary Fig. 3) yields fewer (356) but larger blocks on average (mean ∼60.5; median 22 genes per block) indicative of more continuous macro-synteny despite fewer total blocks. This suggests that collinearity between haplotypes is highly fragmented, reflecting within-genome rearrangements of structural variants (SVs) and copy number variants (CNVs). Overall genome synteny indicates that the genome structure is conserved across alfalfa cultivars grown in different regions, reflecting limited large-scale chromosomal rearrangements during separation.

**Figure 7.**
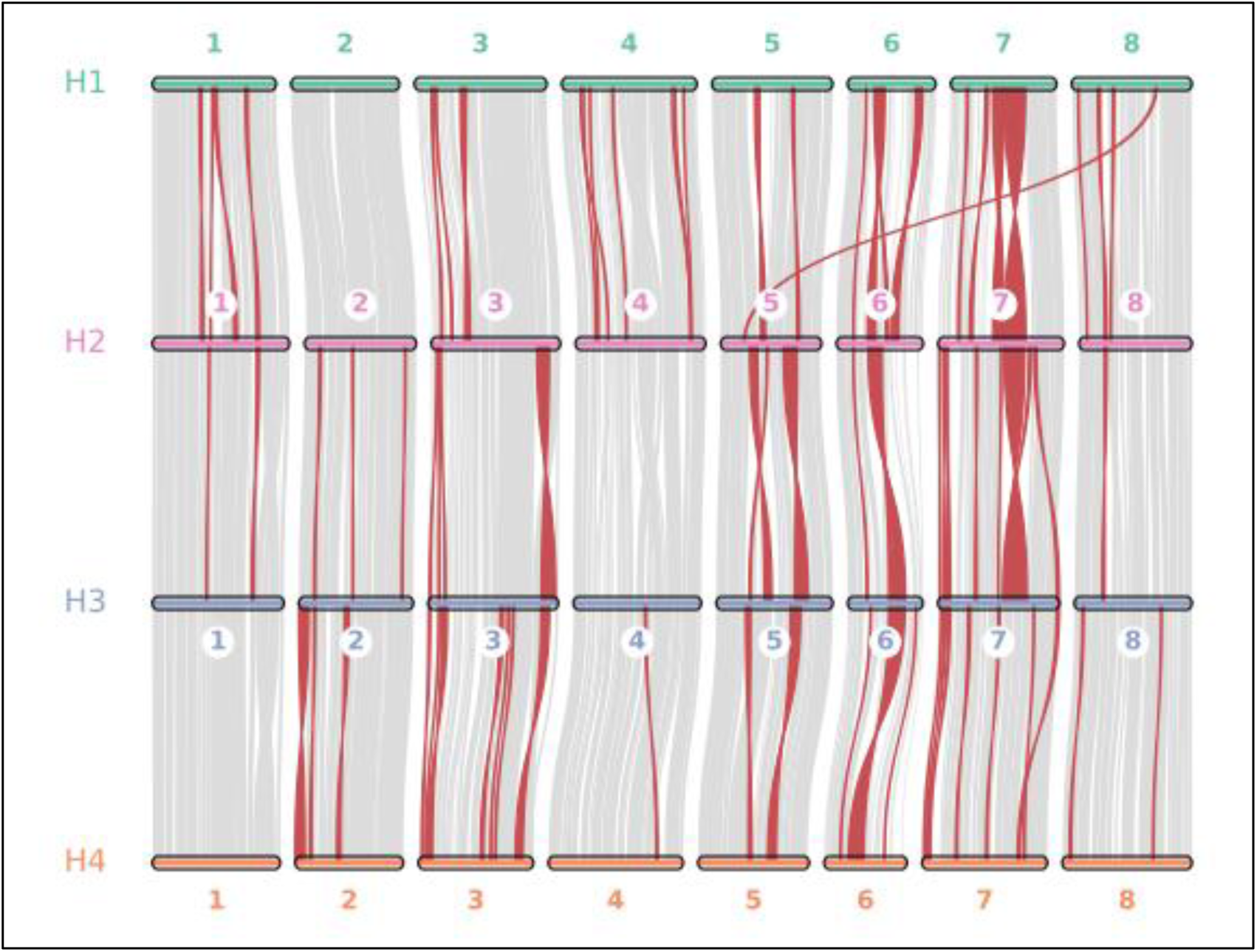
Gene-based synteny among the four haplotypes of the Regen-SY27x genome. The red highlighted syntenic blocks represent inverted syntenic blocks. The figure was generated using the JCVI Python implementation of MCScanX.

Comparative gene family evolution analyses revealed that the Regen-SY27x haplotype-aware genome displays a consistent excess of gene family expansions relative to contractions, whereas Zm1, Zm4, and the XJDY genomes are contraction-biased (Fig. 8). For example, Regen-SY27x haplotype 4 showed 2,978 expanded families compared with only 1,199 contractions, whereas Zm1 exhibited more contractions than expansions (2,912 vs. 1,943). This proliferation of gene families is closely associated with the high Helitron density (5.52%) observed in Regen-SY27x. Specifically, Helitrons were enriched in gene-rich distal chromosome arms (Fig. 1-Track 6), suggesting their proliferation played a pivotal role in driving genomic innovation (Brunner et al. 2005; Hollister and Gaut 2007; Li et al. 2023). This concordance supports the hypothesis that Helitron activity is a key driver of the distinctive evolutionary trajectory of Regen-SY27x compared to other sequenced genotypes.

**Figure 8.**
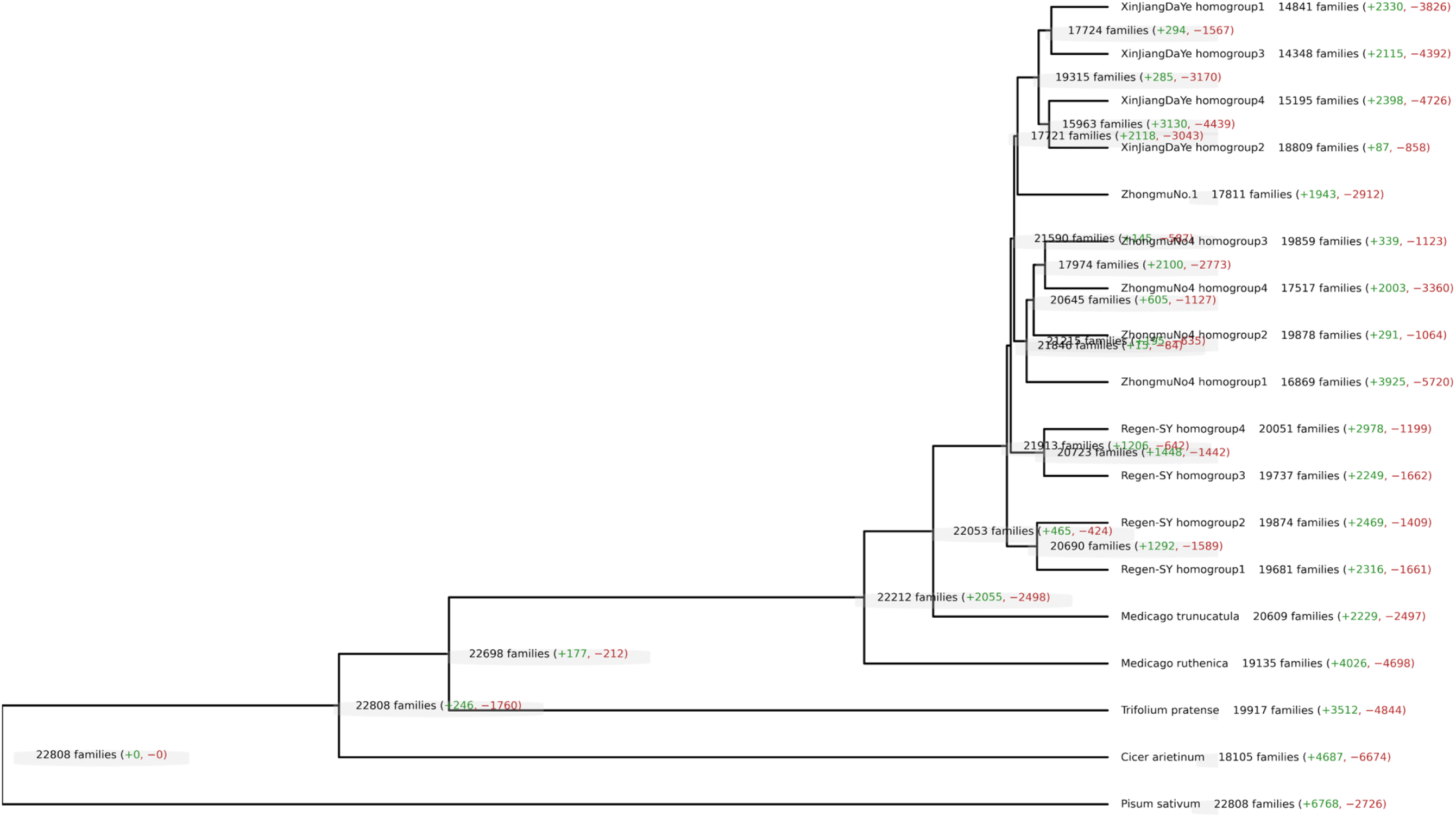
CAFE5 detected gene families with significantly accelerated gene gain or loss on each branch of the phylogenetic tree of different legume species and the sequenced alfalfa genomes Regen-SY27x (four haplogroups), XinJiang DaYe (four haplogroups), ZhongmuNo.1, and ZhongmuNo.4 (four haplogroups).

Across the four haplotypes of the haplotype-resolved Regen-SY27x genome, gene-family turnover was balanced among homologs; each haplotype showed a similar excess of expansions over contractions, and the magnitudes were comparable. In contrast, both XJDY and Zm4 exhibited highly asymmetric turnover between homologs, with some haplotypes being strongly contraction-biased while others showed moderate expansion. These patterns indicate that Regen-SY27x lacks strong haplotype-specific fractionation, whereas XJDY and Zm4 experienced uneven post-polyploid genome remodeling consistent with haplotype-specific gene loss/retention potentially driven by different breeding histories or environmental selection pressures (Shi et al. 2017; Medina et al. 2024).

The gene ontology (GO) enrichment analysis of the top five significantly expanded and contracted gene families across the four Regen-SY27x haplotypes revealed both shared and haplotype-specific functional patterns (Fig. 9). Among expanded genes, terms related to defense response, recognition of pollen, terpene synthase activity, ADP binding, and zinc ion binding were consistently enriched, highlighting the expansion of stress- and defense-related pathways alongside specialized metabolism. In contrast, contracted families were enriched for functions associated with DNA repair, RNA processing, ubiquitin-dependent protein catabolic processes, and telomere maintenance, indicating the loss or reduction of genes linked to genome stability and housekeeping processes. Together, these results suggest that while gene family expansion in alfalfa haplotypes is biased toward functions in defense and environmental adaptation, contraction disproportionately affects genes involved in core cellular maintenance, reflecting contrasting evolutionary pressures shaping the tetraploid genome.

**Figure 9:**
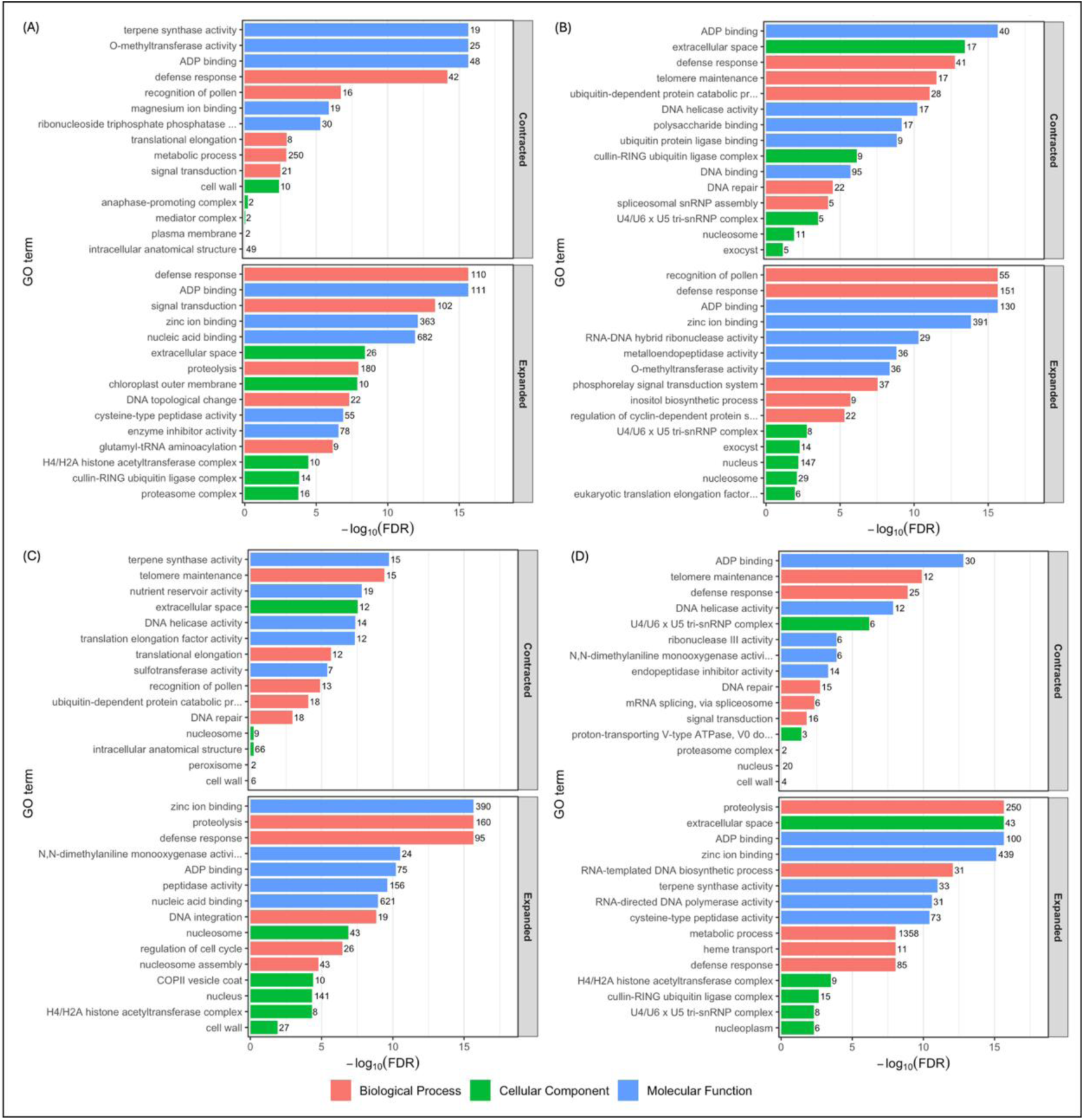
Gene Ontology (GO) enrichment analyses for significantly expanded and contracted genes in the Regen-SY27x genome assembly for (A) haplotype 1, (B) haplotype 2, (C) haplotype 3, (D) haplotype 4. The number on side of bars indicate number of genes for each respective GO term.

### Structural variations

We used Mummer v4.0.0beta2 to align the sequences of pseudochromosome level haplotypes and SyRI to assess the structural variants between them. Our analysis focused exclusively on SVs larger than 100 bp, as larger variants often exert a more profound impact on genome architecture and gene function, while providing higher confidence levels compared to smaller, often more ambiguous, INDELs. Pairwise comparisons among the four Regen-SY27x haplotypes revealed extensive structural divergence, with ∼128,000 to139,000 events identified per haplotype pair (Table 4). The largest categories were duplications and inverted duplications (∼40,000 to 45,000 each), followed by translocations (∼28,000 to 30,000), while insertions and deletions contributed an additional ∼6,000 to 7,000 events each. Inversions were relatively fewer (300 to 340), though spanning a wide size range from <1 kb to >1 Mb. Most SVs were small (<10 kb), indicating that short-range duplications and rearrangements are the predominant form of haplotype variation, likely driven by transposable element activity. The balanced totals across all comparisons suggest that each haplotype contributes a comparable load of structural variants, consistent with high heterozygosity and the synthetic origin of alfalfa cultivars. The dominance of duplication-related SVs implies that gene copy number differences are a major feature of intra-genomic diversity in autotetraploid alfalfa, with potential consequences for traits such as disease resistance and stress adaptation. Comparisons among the four homologs within the Regen-SY27x genome (Table 4, Fig. 10) and, separately, among those in XJDY (Supplementary Table 9, Supplementary Fig. 4) revealed broadly similar SV landscapes. Duplications and inverted duplicates were the most abundant categories (∼40,000 to 46,000 per haplotype pair). Despite these overall similarities, XJDY exhibited consistently higher counts of inversions and translocations compared to Regen-SY27x, indicating greater haplotype reshuffling. Likewise, the slightly elevated total SV counts in XJDY point to modest but notable differences in structural complexity between the two genomes, while still reflecting a shared pattern of extensive haplotype heterogeneity typical of autotetraploid alfalfa.

**Figure 10.**
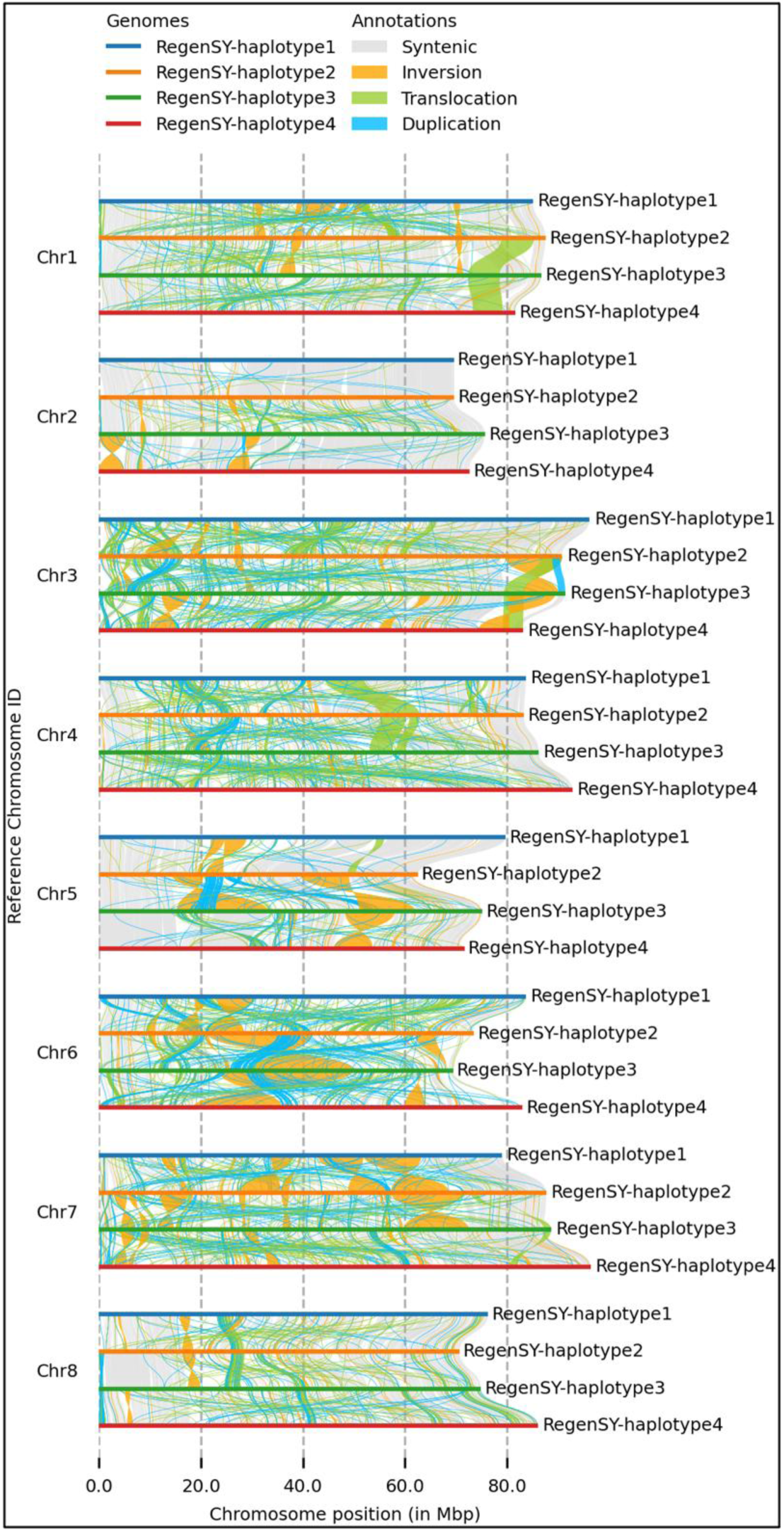
Identification of synteny (gray) and structural rearrangements (inversions, translocations, and duplications) between among the four haplotypes of the Regen-SY27x genome. The figure was generated using the SYRI and plotsr.

**Table 4:** Overview of structural variations identified among Regen-SY27x pseudochromosome level haplotypes.

| Type | Total count | 100 bp to <1 kb | 1 to <10 kb | 10 to <100 kb | 100 to <1 Mb | >1 Mb |
| --- | --- | --- | --- | --- | --- | --- |
| **Regen-SY27x Haplogroup 1 vs Regen-SY27x Haplogroup 2** |  |  |  |  |  |  |
| Inversions | 312 | 102 | 111 | 51 | 28 | 20 |
| Translocations | 28,831 | 14,752 | 12,720 | 1,298 | 55 | 6 |
| Insertions | 5,847 | 4,657 | 983 | 206 | 1 | 0 |
| Deletions | 6,846 | 5,525 | 1,122 | 199 | 0 | 0 |
| Duplicated regions | 38,746 | 17,954 | 19,410 | 1,355 | 26 | 1 |
| Inverted duplicates | 43,023 | 20,629 | 21,049 | 1,327 | 18 | 0 |
| Tandem repeat | 273 | 123 | 99 | 51 | 0 | 0 |
| Copy gain in hap2 | 616 | 437 | 165 | 14 | 0 | 0 |
| Copy lost in hap2 | 3,891 | 2,109 | 1,447 | 328 | 6 | 1 |
| All | 128,385 | 66,288 | 57,106 | 4,829 | 134 | 28 |
| **Regen-SY27x Haplogroup 2 vs Regen-SY27x Haplogroup 3** |  |  |  |  |  |  |
| Inversions | 342 | 123 | 113 | 51 | 39 | 16 |
| Translocations | 30,534 | 15,737 | 13,474 | 1,266 | 51 | 6 |
| Insertions | 6,675 | 5,218 | 1,246 | 210 | 1 | 0 |
| Deletions | 7,606 | 6,169 | 1,238 | 199 | 0 | 0 |
| Duplicated regions | 44,541 | 20,612 | 22,203 | 1,683 | 40 | 3 |
| Inverted duplicates | 43,847 | 20,923 | 21,475 | 1,442 | 7 | 0 |
| Tandem repeat | 280 | 133 | 119 | 28 | 0 | 0 |
| Copy gain in hap3 | 700 | 494 | 197 | 9 | 0 | 0 |
| Copy lost in hap3 | 4,515 | 2,419 | 1,743 | 350 | 3 | 0 |
| All | 139,040 | 71,828 | 61,808 | 5,238 | 141 | 25 |
| **Regen-SY27x Haplogroup 3 vs Regen-SY27x Haplogroup 4** |  |  |  |  |  |  |
| Inversions | 326 | 139 | 86 | 50 | 37 | 14 |
| Translocations | 30,082 | 15,189 | 13,452 | 1,380 | 57 | 4 |
| Insertions | 6,077 | 4,923 | 1,001 | 152 | 1 | 0 |
| Deletions | 6,828 | 5,589 | 1,064 | 174 | 1 | 0 |
| Duplicated regions | 41,882 | 19,321 | 20,944 | 1,579 | 38 | 0 |
| Inverted duplicates | 44,840 | 21,653 | 21,739 | 1,433 | 15 | 0 |
| Tandem repeat | 338 | 140 | 132 | 66 | 0 | 0 |
| Copy gain in hap4 | 722 | 486 | 205 | 31 | 0 | 0 |
| Copy lost in hap4 | 3,992 | 2,189 | 1,476 | 317 | 10 | 0 |
| All | 135,087 | 69,629 | 60,099 | 5,182 | 159 | 18 |

Comparisons of Regen-SY27x with other Chinese autotetraploid cultivars revealed tens of thousands of SVs, underscoring deep divergence among alfalfa germplasms (Table 5). The Regen-SY27x contrast with XJDY yielded ∼187,000 SVs (Supplementary Fig. 5), while Zm4 showed an even higher total of ∼217,000 (Supplementary Fig. 6), both dominated by small insertions and deletions (<10 kb) but also including thousands of large-scale translocations, duplications, and inversions. In contrast, the comparison of Regen-SY27x haplotype 1 with Zm1 identified a much lower SV count (∼58,000), which was expected, as Zm1 is a consensus genome rather than a haplotype-aware assembly. We also identified 8,982,480 SNPs between Regen-SY27x and XJDY as well as 9,599,187 SNPs between Regen-SY27x and Zm4.

**Table 5:** Overview of structural variations identified between Regen-SY27x and other available alfalfa assemblies.

| Type | Total count | 100 bp to <1 kb | 1 to <10 kb | 10 to <100 kb | 100 to <1 Mb | >1 Mb |
| --- | --- | --- | --- | --- | --- | --- |
| **Regen-SY27x vs. XinJiang DaYe** |  |  |  |  |  |  |
| Inversions | 385 | 91 | 56 | 104 | 89 | 45 |
| Translocations | 11,497 | 2,027 | 5,654 | 2,548 | 1,049 | 219 |
| Insertions | 79,852 | 51,610 | 21,524 | 6,685 | 33 | 0 |
| Deletions | 74,397 | 49,316 | 20,395 | 4,673 | 13 | 0 |
| Duplicated regions | 10,790 | 2,618 | 6,307 | 1,587 | 208 | 70 |
| Inverted duplicates | 7,321 | 1,866 | 4,176 | 1,073 | 192 | 14 |
| Tandem repeat | 120 | 83 | 32 | 5 | 0 | 0 |
| Copy gain in XJDY | 476 | 217 | 200 | 58 | 1 | 0 |
| Copy lost in XJDY | 2,361 | 976 | 1,021 | 360 | 3 | 1 |
| All | 187,199 | 108,804 | 59,365 | 17,093 | 1,588 | 349 |
| **Regen-SY27x vs. ZhongmuNo.4** |  |  |  |  |  |  |
| Inversions | 899 | 72 | 128 | 327 | 307 | 65 |
| Translocations | 21,431 | 3,990 | 9,638 | 5,692 | 1,983 | 128 |
| Insertions | 80,302 | 55,496 | 20,226 | 4,558 | 22 | 0 |
| Deletions | 74,724 | 49,735 | 20,354 | 4,627 | 8 | 0 |
| Duplicated regions | 19,268 | 4,224 | 10,440 | 3,864 | 686 | 54 |
| Inverted duplicates | 16,503 | 3,836 | 9,031 | 3,086 | 531 | 19 |
| Tandem repeat | 114 | 61 | 43 | 10 | 0 | 0 |
| Copy gain in Zm4 | 294 | 162 | 102 | 30 | 0 | 0 |
| Copy lost in Zm4 | 3,468 | 1,361 | 1,522 | 576 | 9 | 0 |
| All | 217,003 | 118,937 | 71,484 | 22,770 | 3,546 | 266 |
| **Regen-SY27x Haplotype 1 vs. ZhongmuNo.1** |  |  |  |  |  |  |
| Inversions | 165 | 34 | 28 | 54 | 31 | 18 |
| Translocations | 5,769 | 1,387 | 3,433 | 826 | 119 | 4 |
| Insertions | 20,254 | 13,989 | 5,283 | 982 | 0 | 0 |
| Deletions | 21,076 | 14,540 | 5,379 | 1,157 | 0 | 0 |
| Duplicated regions | 5,006 | 1,264 | 3,196 | 503 | 43 | 0 |
| Inverted duplicates | 4,952 | 1,243 | 3,145 | 512 | 51 | 1 |
| Tandem repeat | 16 | 10 | 3 | 3 | 0 | 0 |
| Copy gain in Zm1 | 107 | 81 | 25 | 1 | 0 | 0 |
| Copy lost in Zm1 | 694 | 358 | 275 | 60 | 1 | 0 |
| All | 58,039 | 32,906 | 20,767 | 4,098 | 245 | 23 |

Other autotetraploid and highly heterozygous species such as potato and sugarcane also exhibit substantial structural and copy number variation, but generally at lower levels of within-genotype haplotype divergence than what we observed in alfalfa. In potato, phased and haplotype-resolved assemblies have revealed numerous SVs, including hemizygosity and tandem CNVs, but the overall scale is lower than the ∼130,000 to 150,000 events we detected among Regen-SY27x haplotypes (Bao et al. 2022; Hoopes et al. 2022; Sun et al. 2022). The structural architecture of the Regen-SY27x genome differs fundamentally from that of the potato cultivars in both scale and composition. While potato structural variation is dominated by large-scale rearrangements including Mb sized inversions that suppress recombination and covers nearly 5% of its gene content (Bao et al. 2022; Sun et al. 2022; Serra Mari et al. 2024; Godec et al. 2025), alfalfa structural variation is characterized by a massive abundance of small-scale events. Specifically, our identification of approximately 40,000 duplicated regions per haplotype comparison, the vast majority of which are smaller than 10 kb (Table 4 and Fig. 9), contrasts with the 220 large-scale duplications (>100 kb) found in potato. This high density of small-scale duplications in alfalfa likely facilitates a more flexible dosage buffer allowing for the maintenance of at least one copy of gene per haplotype (75%) compared to the more fractionated potato genome haplotypes (53.6%). Furthermore, the relative scarcity of megabase-scale inversions in alfalfa may permit higher levels of meiotic recombination, preventing the formation of the large, suppressed-recombination haplotypes that characterize cultivated potato (Sun et al. 2022). Similarly, in sugarcane, an allele-defined genome assembly demonstrated haplotype-specific CNVs and rearrangements, yet again at a different magnitude, scale, and composition compared to the structural divergence in alfalfa (Zhang et al. 2018). These studies highlight that while SVs are common in clonally propagated polyploids, the extraordinarily high intra-genomic SV load in alfalfa is unusual and likely reflects its breeding system that maintains deep haplotype diversity within a single genotype.

Alfalfa cultivars are synthetic populations derived from multiple parents, thereby maintaining high heterozygosity. Simple sequence repeat (SSR) studies have reported high within-variety gene diversity in alfalfa cultivars (Flajoulot et al. 2005; Bagavathiannan et al. 2010). Other studies showed that >80% of variation occurs within cultivars rather than between populations (Nagl et al. 2011; Li et al. 2012). This indicates that genetic diversity has been largely preserved through domestication and breeding, consistent with recent findings that North American core germplasm pools harbor more diversity within individuals than between populations (Medina et al. 2024 Apr). In our study, SV counts between the sequenced genotypes from four different cultivars (187,000 to 217,000) generally exceed those among haplotypes of a single genotype (128,000 to 139,000 in Regen-SY27x). Notably, Regen-SY27x represents U.S. germplasm, whereas the three comparative genomes (XJDY, Zm1, and Zm4) are from Chinese cultivars. The consistently high SV counts in these comparisons reflect not only cultivar-specific breeding histories but also the geographic and genetic divergence between U.S. and Chinese alfalfa germplasm pools, which have largely independent domestication and improvement trajectories. Previous studies have demonstrated pronounced population structure between U.S. and Chinese alfalfa germplasm, with Chinese accessions forming distinct genetic subgroups (Qiang et al. 2015; Long et al. 2022). This geographic separation likely contributes to the deeper structural divergence at genomic level observed between cultivars relative to the already extensive haplotype variation within Regen-SY27x. Additionally, Regen-SY27x is a cross between *M. sativa* ssp. *sativa* and *M. sativa* ssp. *falcata*, two lineages with confirmed genetic differentiation (Bhandari et al. 2011). Population structure analyses have previously confirmed that these subspecies form distinct genetic clusters (Li, Han, et al. 2014), reflecting significant evolutionary distance. This may also contribute to its complex genome structure.

## Conclusion

The haplotype-resolved assembly of Regen-SY27x represents the most complete and structurally informative genome resource to date for autotetraploid alfalfa. The assembly achieves near-chromosome-scale contiguity, balanced haplotype partitioning, and high completeness as evidenced by BUSCO and kmer analyses. Repeat annotation highlights the unusually high abundance of Helitrons and recent bursts of LTR activity, both of which appear to have contributed to extensive gene family expansion, particularly in defense- and stress-related pathways. The NLR repertoire, dominated by TNL and RxNL subclasses and frequently arranged in large tandem clusters, reflects an evolutionarily conserved yet dynamic resistance gene landscape in *Medicago*. Comparative and syntenic analyses reveal strong conservation of chromosomal organization across cultivars while also exposing deep structural divergence among haplotypes and between U.S. and Chinese germplasm pools. Together, these findings underscore the dual nature of the alfalfa genome, conservation at the macro level combined with remarkable intra- and inter-cultivar heterogeneity, driven in part by transposable elements and the species’ outcrossing, synthetic breeding history. This assembly therefore provides a powerful platform for investigating polyploid genome evolution, trait-associated structural variants, and the genetic basis of agronomic improvement in alfalfa. While the current assembly represents a major advancement, there remains room for improvement in phasing and placement of unassigned scaffolds. We, therefore, release this resource as the Regen-SY27x version 1 genome (RSY27v1), providing a foundation for future refinement and community-driven improvement.

## Data Availability

The assembly is available in GenBank under BioProject PRJNA1248037 (BioSample SAMN47849708). The scripts with codes used for identification and annotation of repeat elements, R genes, comparative genomics are provided in the Github repository https://github.com/shannonlabumn/Regen-SY27xGenome. Scripts for adjudicating the gene annotation are in https://github.com/ncgr/Adjudicator. Plant material is available on request.

## Acknowledgements

We thank Melinda Dornbusch for plant maintenance. Mention of any trade names or commercial products in this article is solely for the purpose of providing specific information and does not imply recommendation or endorsement by the U.S. Department of Agriculture. USDA is an equal opportunity provider and employer.

